# AA-ending codon-based computational analyses uncover novel cytoplasmic effectors in the *Magnaporthe oryzae* secretome

**DOI:** 10.1101/2025.11.10.687658

**Authors:** Gang Li, Sudeep Tiwari, Nawaraj Dulal, Ben Wheeler, Richard A. Wilson

## Abstract

Phytopathogens secrete diverse, sequence-unrelated effector proteins outside (apoplastic) or into (cytoplasmic) plant cells to suppress host defenses and cause devastating diseases. Effectors rapidly evolve to avoid counter-detection by the host, and fungal cytoplasmic effectors lack recognizable host cell-targeting motifs, impeding cataloguing of the full suite of host-deployed effectors encoded in pathogen genomes. In the devastating rice blast fungus *Magnaporthe oryzae*, cytoplasmic and apoplastic effectors are secreted by different routes, but candidates must be confirmed by fluorescent labelling, consequently few are known. AA-ending codon usage rates are the fraction of AA-ending codons as a percentage of the total number of AA- and synonymous AG-ending codons in an mRNA. In *M. oryzae*, high AA % rates of the corresponding mRNAs distinguished seven experimentally confirmed cytoplasmic effectors from four experimentally confirmed apoplastic effectors, but whether AA % rates can identify new effectors is unknown. Here, using computational analyses and live-cell imaging confirmation, we successfully predicted, based on AA % rates, two new cytoplasmic effectors and two apoplastic effectors in the *M. oryzae* secretome. Our findings support the notion that cytoplasmic effector mRNAs are globally enriched for AA-ending codons, aiding effector discovery and the search for novel sources of durable crop resistance.

## Introduction

Biotic plant diseases are intractable threats to global food security. To colonize host plant tissue, fungal plant pathogens secrete two classes of virulence factors to promote infection: apoplastic effectors are secreted into the matrix between host plasma membrane and pathogen cell wall to (for example) prevent host immune system detection of fungal cell wall components; cytoplasmic effectors are secreted into the host plant cell to manipulate cellular processes and suppress plant innate immunity (Valent, 2025). In a classic evolutionary arms race, intracellular host *R* gene products (directly or indirectly) recognize cognate cytoplasmic effectors (which are then termed avirulence (*AVR*) gene products), thereby triggering immunity (Jones and Dangl, 2006). However, pathogens can lose or rapidly mutate *AVR* genes to overcome host *R* gene product detection, resulting in host jumps and new epidemics (Valent, 2025).

Identifying cytoplasmic effectors and understanding effector evolution will help safeguard agriculture by determining which *R* genes to deploy against which pathogen populations, by finding new cognate *R* genes and, potentially, by aiding the design of bespoke R proteins when suitable host *R* genes are unavailable (Seong and Krasileva, 2021; Wilson and McDowell, 2022; Kourelis et al., 2023; Valent, 2025). It is unfortunate, then, that progress towards identifying new fungal cytoplasmic effectors is impeded due to the difficulty in predicting them from first principles in the secretome. They are highly diverse at the nucleotide level and lack recognizable secretion-associated motifs besides the N-terminal signal peptide (SP) sequences carried by all secreted proteins; thus, compared to oomycete cytoplasmic effectors that carry RxLR motifs, few fungal host-deployed effectors are known (Sperschneider and Dodds, 2022; Wilson and McDowell, 2022).

The devastating rice blast fungus *Magnaporthe oryzae* (syn. *Pyricularia oryzae*), which causes 10-30 % of global rice losses annually, grows biotrophically in living rice cells for the first few days of infection, secreting effectors to avoid detection (Wilson and Talbot, 2009; Wilson 2021; Valent, 2025). During biotrophy, *M. oryzae* secretes SP-carrying apoplastic effectors via the brefeldin A (BFA)-sensitive conventional ER-Golgi secretion pathway (Giraldo et al., 2013). In contrast, SP-carrying cytoplasmic effectors are secreted into the plant-lipid rich biotrophic interfacial complex (BIC) via a BFA-resistant Type IV unconventional protein secretion (UPS) pathway that bypasses the Golgi (**Fig. 1a**) (Giraldo et al., 2013; Li et al., 2023a; Dulal and Wilson, 2024). From the BIC, cytoplasmic effectors are translocated into host cells via clathrin-mediated endocytosis (Oliveira-Garcia et al., 2023). Unconventional secretion and accumulation in the BIC are hallmarks of *M. oryzae* cytoplasmic effectors, but cytoplasmic and apoplastic effectors cannot be distinguished in the genome. Consequently, although *M. oryzae* cytoplasmic effectors can be inferred genetically as *AVR* genes that trigger immunity in the presence of host cognate *R* genes, they must be verified empirically on a case-by-case basis using fluorescent tags, limiting discovery (Li et al., 2023a).

**Figure 1.**
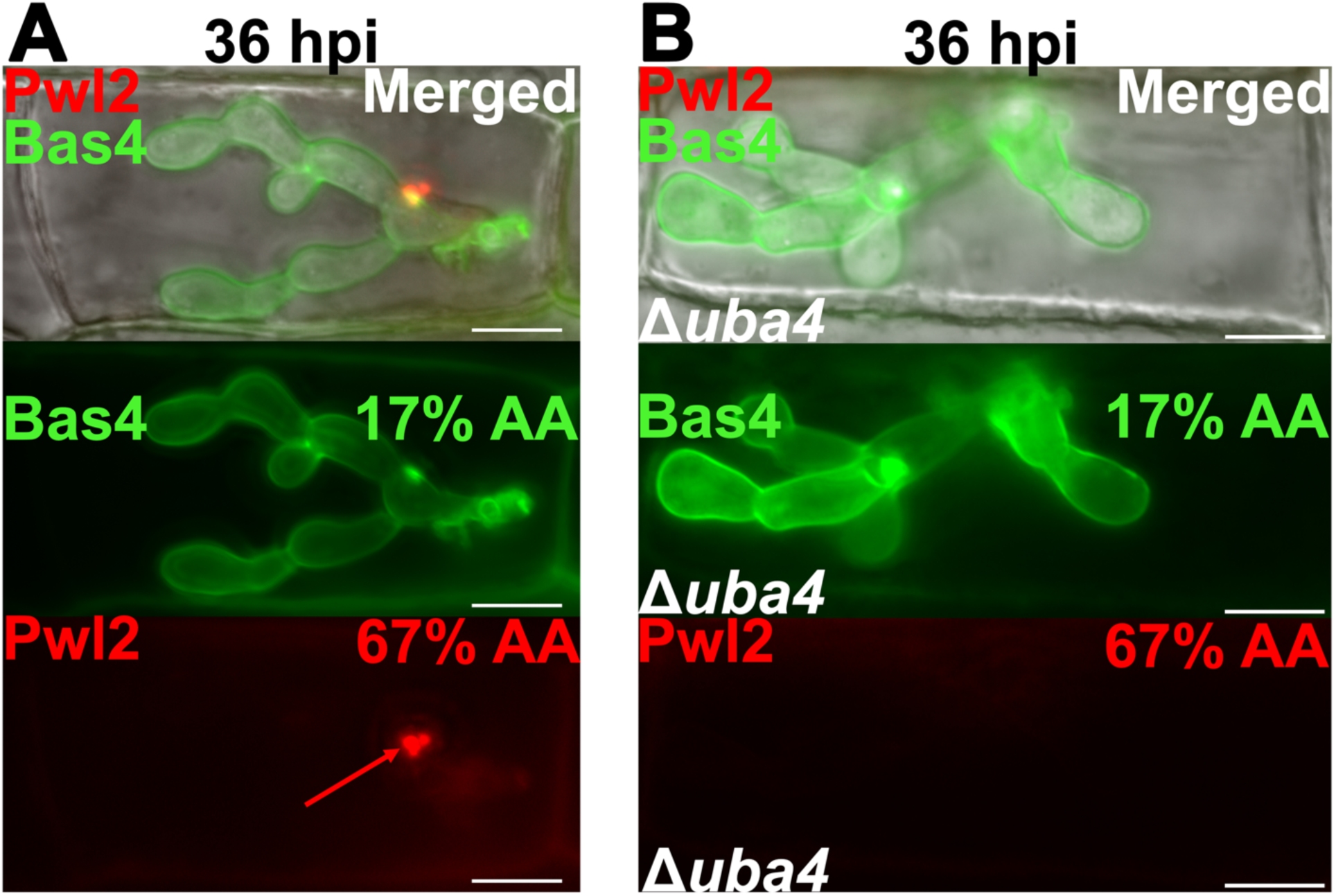
Cytoplasmic effectors like Pwl2 accumulate in the BIC while apoplastic effectors like Bas4 localize in the apoplastic compartment. **A**. During biotrophy, confirmed cytoplasmic effectors like Pwl2 are secreted by the Type IV Golgi bypass unconventional protein secretion pathway (Giraldo et al., 2013; Dulal and Wilson, 2024) into the biotrophic interfacial complex (BIC), where they accumulate before translocating into host cells. Apoplastic effectors like Bas4 are secreted into the apoplastic space by the conventional ER-Golgi secretion route (Giraldo et al., 2013; Dulal and Wilson, 2024). Live-cell imaging shows the biotrophic growth in a living host rice cell of the WT rice-infecting isolate Guy11 carrying the pBV591 vector (Khang et al., 2010) and expressing *PWL2-mCherry:NLS* and *BAS4-eGFP* under native promoters. B. *PWL2* mRNA translation but not *BAS4* mRNA translation is dependent on tRNA thiolation mediated by Uba4 (Li et al., 2023). A,B. Images were taken at 36 hr post inoculation (hpi). Red arrow indicates Pwl2 accumulation in the BIC. Bars are 10 µm. % value refers to the AA % rate for the indicated effector mRNA (Li et al., 2023a). Images are representative of 50 infected rice cell observations per leaf sheath, repeated in triplicate.

Clues as to how cytoplasmic and apoplastic effectors may be distinguished in both the secretome and by the cell during secretion come from our recent findings showing that in *M. oryzae*, the efficient decoding of AA-ending codons in effector mRNAs is essential for the production and secretion of at least a small set of previously confirmed cytoplasmic but not apoplastic effectors (Li et al., 2023a). AA-and synonymous AG-ending codons encode lysine (AAA and AAG), glutamine (CAA and CAG) and glutamic acid (GAA and GAG). In fungi, efficient decoding of AA-ending codons (but not synonymous AG-ending codons) requires the Uba4-Urm1 sulfur relay system, which thiolates the anticodon wobble uridine at position 64 (s^2^U_34_) of cognate tRNAs (Rezgui et al., 2013; Nedialkova and Leidel, 2015). The s^2^U_34_ modification is universal and its loss leads to ribosomal pausing at AA-ending codons (Rezgui et al., 2013; Nedialkova and Leidel, 2015; Li et al., 2023a; Zhang et al., 2025). In *M.* oryzae, loss of the s^2^ modification in Δ*uba4*- and *Δurm1*-carrying mutant strains abolished the translation of *PWL2*, *AVR-Pita* and *AVR-Pik* cytoplasmic effector mRNAs enriched for AA-ending codons, but did not affect the production and conventional secretion of the Bas4 and Slp1 apoplastic effectors, whose mRNAs are enriched for synonymous AG-ending codons instead (Li et al., 2023a). In the Δ*uba4* mutant strain, recoding all AA-ending codons in *PWL2* to synonymous AG-ending codons restored Pwl2 production and secretion (Li et al., 2023a). Expressing this recoded version of *PWL2* in wild type resulted in unregulated super-secretion of Pwl2 into the BIC, which expanded and fractured, resulting in the loss of virulence (Li et al., 2023a). Thus, AA-ending codon usage optimizes cytoplasmic effector translation and secretion to maintain BIC integrity. This raises the question of whether AA-ending codon enrichment in cytoplasmic effector mRNAs is global and therefore sufficient to distinguish cytoplasmic effectors from apoplastic effectors in the secretome. If so, this would be a powerful tool for identifying novel effectors from first principles.

This study was motivated by asking, can AA-ending codon-enrichment predict novel cytoplasmic effectors in the *M. oryzae* secretome? This is important because although the *M. oryzae* secretome comprises 1854 genes encoding proteins less than 1000 amino acids in length and therefore likely to be effectors (Seong and Krasileva, 2021), how many are cytoplasmic effectors, how many are apoplastic effectors, and indeed how many are small, secreted enzymes, is unknown, hindering effector discovery and understanding. Here, using computational approaches and live cell-imaging, we correctly identified and confirmed two new cytoplasmic effectors based on the high AA-ending codon usage rates of their associated mRNAs. Transcriptomic analysis showed that, like for other known cytoplasmic effector mRNAs, these were expressed at very low levels *in planta* during biotrophic growth. Conversely, we found one new apoplastic effector gene in the secretome based on its high *in planta* expression level and low AA-ending codon content. We also correctly established that a putative effector with no AA-ending codons and 16-AG ending codons in the corresponding mRNA, previously predicted by *in silico* molecular docking modelling to be a cytoplasmic effector, was in fact localized to the apoplast, thus directly demonstrating the superior power of AA-ending codon-enrichment for predicting cytoplasmic effectors compared to other computational approaches. Together, our findings indicate that cytoplasmic effector mRNAs accumulate AA-ending codons in a non-neutral manner, allowing them to be distinguished from apoplastic mRNAs. By augmenting effector discovery, this work paves the way to a deeper understanding of effector evolution and secretion that may lead to new sources of durable crop resistance.

## Results

### AA-ending codons accumulate in *M. oryzae* cytoplasmic effector mRNAs across isolates and host-infecting lineages

Using AA-ending codon usage rates (AA % rates) (the fraction of AA-ending codons as a percentage of the total number of AA-ending and synonymous AG-ending codons in an mRNA), we first sought to understand how widespread, and thus how robust, was the rule that cytoplasmic (but not apoplastic) effector mRNAs carry more AA-ending codons than synonymous AG-ending codons. Previously, from a small pool of four apoplastic and seven cytoplasmic effectors with stringent localization and secretion route support (Li et al., 2023a), we had further demonstrated that three of these cytoplasmic effector mRNAs – *PWL2*, *AVR-Pik* and *AVR-Pita* with AA % rates of 67 %, 92 % and 88 %, respectively – required Uba4 for translation, but two apoplastic effector mRNAs – *BAS4* (17 % AA) and *SLP1* (7 % AA) – did not (shown in **Fig. 1B** for *PWL2-mCherry:NLS* and *BAS4-eGFP* expressed under their own promoters from the pBV591 vector) (Li et al., 2023a). By comparison, calculating the AA % rate for all protein codon sequences in the *M. oryzae* genome yielded a value of 29 % (Li et al., 2023a). Thus, at least a subset of cytoplasmic effector mRNAs is enriched for AA-ending codons over synonymous AG-ending codons, whereas apoplastic effector mRNAs, and the *M. oryzae* genome in general (like for human and mice genomes (Goffena et al., 2018)), is depleted for AA-ending codons in favor of synonymous AG-ending codons. This AA-ending codon usage bias is reflected in the reduced number of putative cognate tRNAs matching AA-ending codons (12 genes) compared to tRNAs matching synonymous AG-ending codons (39 genes) in the *M. oryzae* 70-15 reference strain, according to tRNAscan-SE (Chan et al., 2021). The concentration of each tRNA in the cell is determined by the number of gene copies in the genome (Tuller et al., 2010) and impacts codon translation efficiency because the most abundant codons pair with the most abundant tRNAs (Novoa and Pouplana, 2012; Novoa et al., 2012). Therefore, AA-ending codons are likely slowly translated (even in WT) and are considered non-optimal because they are both rare in the genome and have a low number of cognate tRNAs, whereas AG-ending codons are considered optimal because their high abundance in the genome matches the high abundance of cognate tRNA genes, ensuring they are rapidly decoded (Plotkin and Kudla, 2011; Chaney and Clark, 2015).

We use the *M. oryzae* rice-infecting WT isolate Guy11 for physiological studies, and the reference genome sequence from the rice-infecting laboratory strain 70-15, which is derived from Guy11 (Dean et al., 2005), for molecular genetics studies. In addition to rice, the *M. oryzae* species complex infects a wide-range of grasses with host species-specificity determined by avirulence genes in a classic gene-for-gene interaction (Valent, 2025). We expanded our analysis of AA-ending codon usage in *M. oryzae* effectors by examining putative cytoplasmic effectors from different *M. oryzae* host-infecting lineages that were either homologues of known effectors, or that had been inferred by genetics to be avirulence factors (and therefore likely host-targeted to interact with the corresponding host *R* gene product), but which may lack experimental support for their deployment. We first assessed AA % rates for orthologues of the cytoplasmic effector-encoding *PWL2* gene, a highly prevalent host-specificity gene (Valent, 2025). The 70-15 reference genome has two copies of *PWL2*, one encoded at the locus MGG_13863, which is expressed in our studies in Guy11 under its native promoter from the vector pBV591 (**Fig. 1A**), and one at the MGG_04301 locus. Both mRNA sequences comprise 14 AA- and 7 AG-ending codons from a total of 145 codons, yielding identical AA % rates of 67 % (**Table 1**). In addition, *PWL2* sequences from *M. oryzae* isolates causing blast on crab grass, foxtail millet and finger millet, and from the *M. oryzae* relative *Pyricularia grisea* (Gómez Luciano et al., 2019), have identical or near-identical sequences to MGG_13863 (**Table 1**). Thus, elevated AA % rates are conserved in *PWL2* sequences from across a wide range of geographically distinct *M. oryzae* host-specific isolates and at least one related species.

**Table 1.** AA % rates of effector mRNAs from across isolates, host-infecting lineages and related species.

| Gene | GenBank Accession | AA % rate | Isolate | Host | Species | Geographical location |
| --- | --- | --- | --- | --- | --- | --- |
| *PWL2* | MGG_13863 | 67 | 70-15 | Rice | *M. oryzae* | Laboratory strain |
| *PWL2* | MGG_04301 | 67 | 70-15 | Rice | *M. oryzae* | Laboratory strain |
| *PWL2* | MN072513 | 64 | BJM-1PWL2 | Crab grass | *M. oryzae* | China |
| *PWL2* | MN072505 | 67 | HuBG-1 | Bristle grass | *M. oryzae* | China |
| *PWL2* | MT669815 | 67 | E22 | Finger millet | *M. oryzae* | Ethiopia |
| *PWL2* | PgNI_12163 | 64 | NI907 | Crab grass | *P. grisea* | Japan:Tochigi |
| *PWL2* | PgNI_02586 | 64 | NI907 | Crab grass | *P. grisea* | Japan:Tochigi |
| *PWL1* | MT669814 | 73 | E22 | Finger millet | *M. oryzae* | Ethiopia |
| *PWT3* | LC202650.1 | 55 | Br58 | Oat | *M. oryzae* | Brazil |
| *PWT4* | LC202655 | 82 | Br58 | Oat | *M. oryzae* | Brazil |
| *AvrPi54* | HF545677 | 0 | Mo-nwi-55 | Rice | *M. oryzae* | India |

Along with *PWL2*, the Ethiopian finger millet blast isolate E22 also carries *PWL1*, a well-studied *AVR* gene preventing pathogenicity on weeping lovegrass that is absent from the rice-infecting 70-15 and Guy11 genomes (Khang et al., 2010; Masaki et al., 2023). *PWL1* from the E22 isolate was identical to sequences from 49 Ethiopian, 35 Tanzanian, and two Kenyan finger millet blast strains (Masaki et al., 2023) as well as to *PWL1* from a previous isolate whose protein was shown to be BIC localized (Khang et al., 2010). All versions of *PWL1* have a high AA % rate of 73 % (16 AA- and 6 AG-ending codons in the 147-codon mRNA) (**Table 1**). We hypothesized that this high AA-ending codon-enriched mRNA from a different host-infecting lineage was, like *PWL2* from rice-infecting isolates, translationally controlled by Uba4 to produce an unconventionally secreted cytoplasmic effector. To test this, we transformed Guy11 WT and Δ*uba4* protoplasts with a modified pBV591 vector expressing *PWL1* under its native promoter and fused to *mCherry:NLS* (along with *BAS4* under its native promoter fused to *eGFP*), and inoculated the resulting strains onto optically clear detached rice leaf sheaths from 3-week-old seedlings of the susceptible cultivar CO-39. For each strain, at least three independent transformants expressing *PWL1-mCherry:NLS* and *BAS4-GFP* were examined and found to be identical. In WT, by 36 hpi, Pwl1 was secreted into the BIC in a BFA-insensitive manner, but Pwl1 production was not observed in the Δ*uba4* mutant unless infected leaf sheaths were treated with paromomycin (**Fig. 2A**), which increases near-cognate tRNA acceptance in the ribosome and was shown previously to restore *PWL2*, *AVR-Pita* and *AVR-Pik* mRNA translation in Δ*uba4* (Li et al., 2023a). Thus, the host-specific effector Pwl1 from the finger millet isolate E22 is, like Pwl2 from the rice isolate 70-15, unconventionally secreted into the BIC in a manner requiring Uba4-dependent efficient decoding of its AA-ending codon-enriched mRNA.

**Figure 2.**
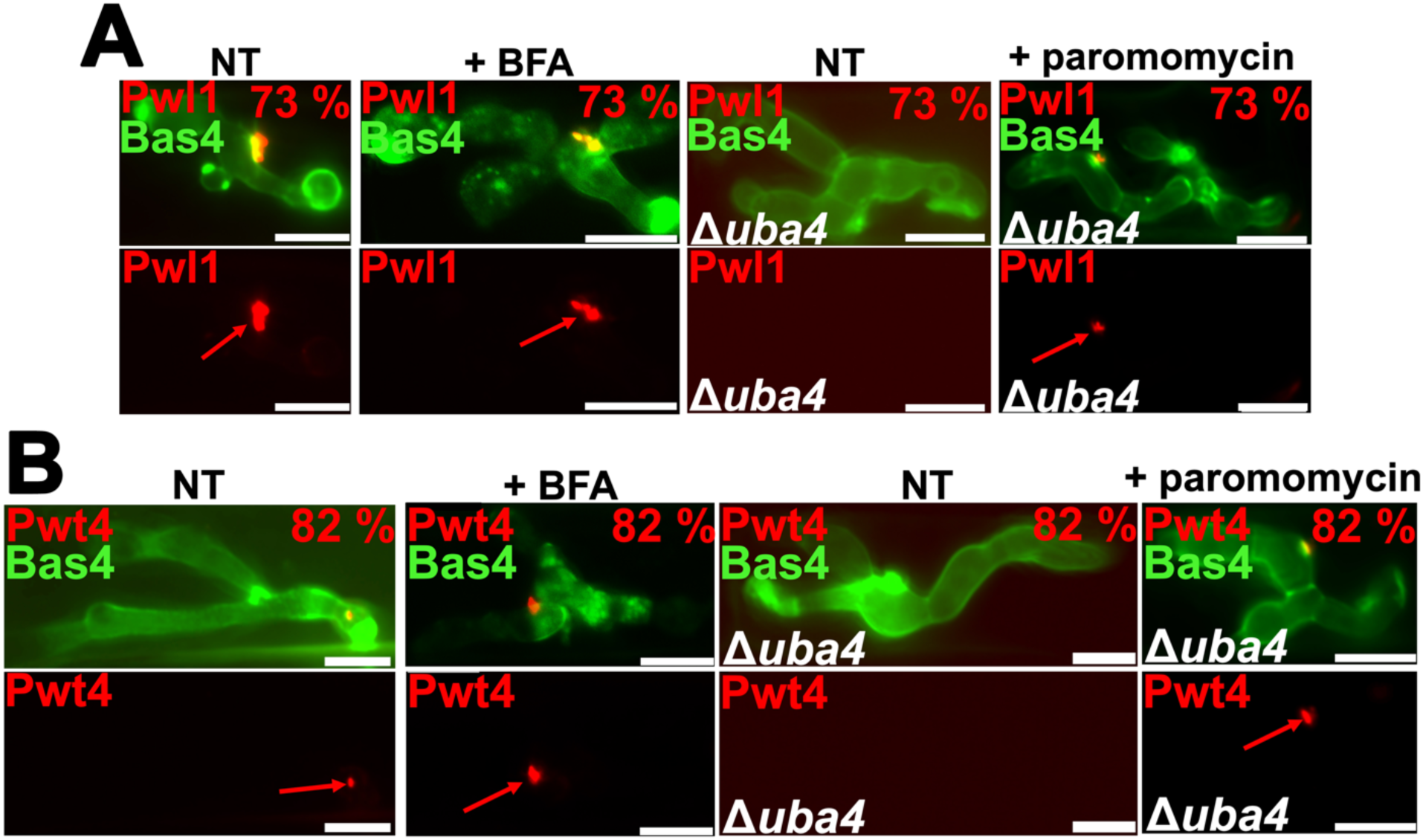
*PWL1* and *PWT4* -important for finger millet blast and wheat blast, respectively - have high AA % rates and are translated and secreted unconventionally in a Uba4-dependent manner. **A,B.** Live-cell images of the indicated strains and treatments were taken at 36 hpi for the non-treated (NT) controls. For BFA treatments, samples were treated at 32 hpi and imaged at 37 hpi. For paromomycin treatments, samples were treated at 36 hpi and imaged at 44 hpi. Red arrows indicate effector accumulation in the BIC. Bars are 10 µm. % value refers to the AA % rate for the indicated effector mRNA (Li et al., 2023a). Images are representative of 50 infected rice cell observations per leaf sheath, repeated in triplicate.

Wheat blast caused by host adapted *M. oryzae* strains is emerging as a prominent threat (Valent, 2025). *PWT3* and *PWT4* from the *Avena* isolate Br58 are host species specificity genes which condition its avirulence on wheat (Inoue et al., 2017). *Lolium* isolates also carry copies of these genes. *Avena* and *Lolium* blast isolates are incompatible on wheat due to the presence of wheat *R* genes corresponding to *PWT3* and *PWT4*. Loss of the corresponding *R* genes, or loss of *PWT3* and/ or *PWT4*, leads to virulence on wheat (Inoue et al., 2017). The AA % rate for *PWT3* is 55 % (12 AA-ending codons and 10 AG-ending codons in 141 codons) and for *PWT4* it is 82 % (14 AA- and 3 AG-ending codons out of 282 codons) (**Table 1**). We transformed a modified pBV591 vector expressing the *Avena* isolate *PWT4* gene under its native promoter fused to *mCherry:NLS*, along with *BAS4-eGFP* under its native promoter, into Guy11 and Δ*uba4* protoplasts, finding that, by 36 hpi, Pwt4 was unconventionally secreted into the BIC in a Uba4-dependent manner (**Fig. 2B**). *PWT4* mRNA translation was restored in Δ*uba4* by paromomycin treatment (**Fig. 2B**). Thus, translation of the AA-ending codon-enriched *PWT4* mRNA – like *PWL1* and the previously reported *PWL2*, *AVR-Pita* and *AVR-Pik* mRNAs (Li et al., 2023a) – requires efficient AA-ending codon decoding by the Uba4-Urm1 sulfur relay system. In addition to bolstering support for the notion that cytoplasmic effector mRNAs are globally enriched for AA-ending codons, by showing for the first time that Pwt4 is unconventionally secreted and BIC-localized, our results provide new information on how these important virulence determinants are deployed.

We conclude that known or inferred cytoplasmic effector mRNAs from across *M. oryzae* isolates, host-infecting lineages and a closely related species have high AA % rates, suggesting AA-ending codon accumulation is a globally conserved feature of cytoplasmic but not apoplastic effector mRNAs. At least five AA-ending codon-enriched cytoplasmic effector mRNAs are now known to require the Uba4-Urm1 sulfur relay system for translation.

### Reclassification of a predicted cytoplasmic effector to a confirmed apoplastic effector based on an AA % rate of 0

*AVR* gene products interact with host intracellular receptors and should therefore be cytoplasmic effectors. Some *M. oryzae AVRs* from rice-infecting isolates are characterized genetically and have known corresponding host *R* genes, but the Avr proteins have no localization and/ or secretion route support. We calculated the AA % rates for the *M. oryzae* rice isolate *AVR* genes whose proteins were not included in our original, stringent analysis (Li et al., 2023a): *Avr-Pii* (75 % AA), *AvrPib* (64 %), *AvrPi9* (encoding a BIC-localized protein with unconfirmed secretion route (Wu et al., 2015)) (38 %), and *AvrPi54* (0 % AA). *AvrPi54* (**Table 1**) was identified by *in silico* molecular docking modelling in the genome of the *M. oryzae* rice isolate Mo-nwi-55 as encoding a putative interactor of the protein encoded by the *R* gene *Pi54* (Ray et al., 2016). The 153 codon *AvrPi54* gene (which is absent from Guy11 and 70-15) carries no AA-ending codons and 16 AG-ending codons and may refute our hypothesis that cytoplasmic effector mRNAs are globally enriched for AA-ending codons. Alternatively, the AvrPi54 protein may instead be apoplastic and interact with its cognate *R* gene product by indirect or currently unknown means. To assess this, we expressed in Guy11 the *AvrPi54* gene fused to *mCherry:NLS*. Because the *AvrPi54* native promoter has not been experimentally delineated, expression was achieved using the *PWL2* promoter (Khang et al., 2010). We found that AvrPi54 localized to the apoplast in a Uba4-independent manner (**Fig. 3**). Thus, in this case, our AA % rule holds true, and the 0 % AA rate of *AvrPi54* mRNA correctly predicts it encodes an apoplast-localized effector. Incidentally, these results also show that the *PWL2* promoter alone is not sufficient to determine protein localization.

**Figure 3.**
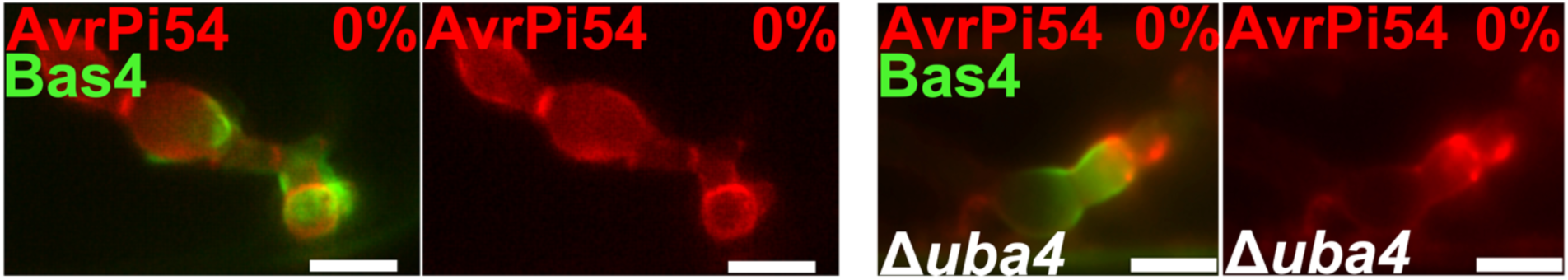
*AVR-Pi54* mRNA does not require Uba4 for translation and encodes an apoplastic effector. Live-cell images of the indicated strain were taken at 36 hpi. Bars are 10 µm. % value refers to the AA % rate for the indicated effector mRNA (Li et al., 2023a). Images are representative of 50 infected rice cell observations per leaf sheath, repeated in triplicate.

### mRNAs with high AA-ending codon usage rates are rare in the *M. oryzae* secretome but correlate with small mature protein size

The previous *AVR-Pi54* results showed AA % rates outperformed structural *in silico* approaches in identifying cytoplasmic effectors, prompting us to ask whether AA-ending codon usage could predict new effectors. To address this, we first sought to determine the prevalence of AA-ending codon-enriched genes in the *M. oryzae* secretome. Both apoplastic and cytoplasmic effectors, like secreted enzymes and membrane proteins, have recognizable N-terminal SPs targeting them to the ER. SPs direct translating cytosolic ribosomes to the ER, where they become membrane-bound, allowing nascent proteins to be co-translationally translocated into the ER lumen before SPs are cleaved. Recent analyses using the neural network of SignalP v3.0 followed by TMHMM v2.0 to eliminate putative integral membrane proteins, and then the removal of mature proteins with ≥ 1000 amino acids, identified 1854 likely secreted proteins encoded in the *M. oryzae* 70-15 reference genome (Seong and Krasileva, 2021). We calculated the AA % rates for each protein coding sequence in this dataset and ordered the genes based on their AA % rates (**Fig. 4A**; **Supplementary Table S4**). The AA-ending codon usage rate for the entire secretome was 23.15 %, lower than the 29 % calculated for the genome. Only 13 % (244) of secretome genes have ≥ 50 % AA-ending codon usage rates (by comparison, 4 % of secretome genes have 0 % AA rates) (**Fig. 2A**; **Supplementary Table S4**), suggesting AA-ending codon-enriched mRNAs are rare in the secretome. Two genes, MGG_15126 and MGG_16011, have 100 % AA rates, but their rates are based on 2 and 1 AA-ending codon(s), respectively, and no AG-ending codons (**Supplementary Table S4**), therefore we did not consider that they were AA-ending codon-enriched. The 70-15 *AVR-PikD* allele at MGG_15972 (Seong and Krasileva, 2021) has the fourth highest AA % rate of 92.31 % derived from 15 AA- and 1 AG-ending codon(s) in 113 codons; *AVR-Pita* (MGG_15370) has the seventh highest AA % rate of 87.88 % from 29 AA- and 4 AG-ending codons in 224 codons (**Fig. 4A**). The other genes at the top positions in **Supplementary Table S4** were uncharacterized.

**Figure 4.**
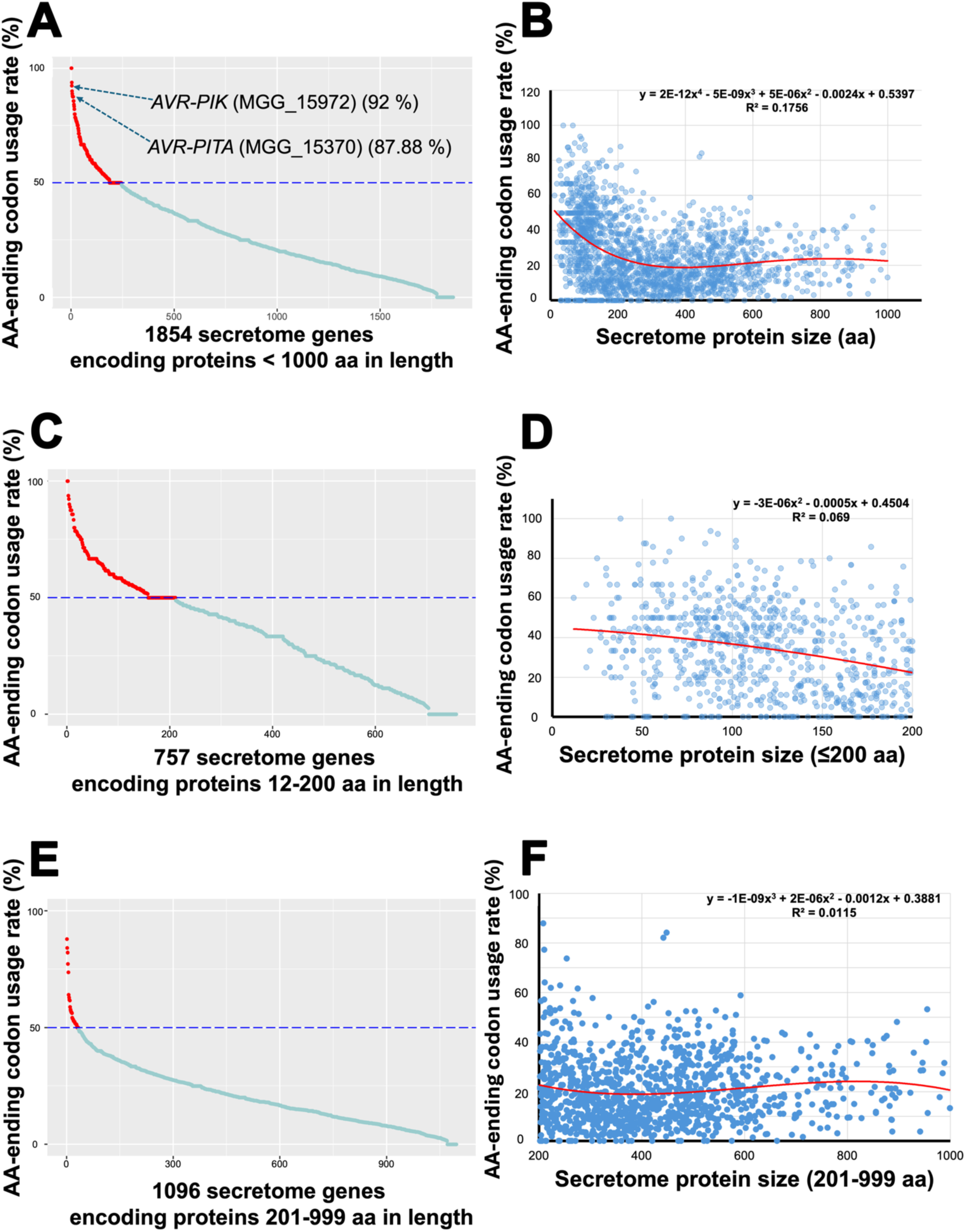
Rare AA-ending codons accumulate in mRNAs encoding small, secreted proteins. A. 13. % of secretome genes (244 out of 1854) have AA % rates ≥ 50 %. The AA-ending codon usage rates of the 1854 genes in the *M. oryzae* secretome dataset were determined using coRdon (Elek et al., 2024). **B**. AA % rates tend to decrease in secretome mRNAs as the encoded mature protein size (in amino acids) increases. The x-axis shows secretome genes encoding proteins of the indicated size (ranging from 12 to 999 amino acids), the y-axis represents the AA % rate of the corresponding mRNA. A red fourth-order polynomial regression curve is overlaid on the scatter graph, with the equation and R² value displayed. Spearman’s rank correlation analysis revealed a statistically significant, moderate negative correlation between protein size and AA-ending codon rate (*ρ* = -0.311, two-tailed *p* < 0.001; 99% CI: –0.3651 to – 0.2538). **C**. 28 % (211) of the 758 secretome genes encoding mature proteins with ≤ 200 amino acids have AA % rates ≥ 50 %. **D**. For the 757 secretome genes encoding mature proteins with ≤ 200 amino acids, AA % rates tend to decrease with increasing mature protein size. A second-order polynomial regression curve (red) is overlaid on the scatter graph, with the equation and R² value shown on the plot. Spearman’s rank correlation analysis revealed a statistically significant, moderate negative correlation between protein size and AA-ending codon rate (*ρ* = -0.2590, two-tailed *p* < 0.001; 99% CI: –0.3466 to – 0.1670). **E**. 3 % (33) of the 1096 secretome genes encoding mature proteins with 201-999 amino acids have AA % rates ≥ 50 %. **F**. For the 1096 secretome genes encoding mature proteins with > 200 amino acids, AA % rates were not correlated with mature protein size. A red third-order polynomial regression curve is overlaid on the scatter graph, with its equation and R² value displayed. Spearman’s rank correlation analysis revealed no statistically significant correlation between protein size and AA-ending codon usage (*ρ* = 0.07054, two-tailed *p* = 0.02; 99% CI: –0.009560 to 0.1497). **A,C,E**. Genes were ranked in descending order based on the AA % rate of each gene. The blue dashed line indicates 50 % AA rate. Red dots are genes with ≥ 50 % AA rates, green dots are genes with < 50 % AA rates. **B,D,F**. Charts were generated using Microsoft Excel.

The distribution of AA % rates in the secretome is continuous (**Fig. 4A**) and does not form two distinct groups of data points that might represent cytoplasmic and apoplastic effector mRNAs. To better understand the relationship between AA-ending codon usage rates and the secretome, we analyzed the correlation between AA % rates and mature protein size (ie. after the SP has been removed). For all 1854 secretome genes, encoding mature proteins ranging in size from 12 – 999 amino acids, Spearman’s rank correlation analysis revealed a statistically significant, moderate negative correlation between mRNA AA-ending codon usage rates and the size of their corresponding mature proteins (*ρ* = -0.311, two-tailed *p* < 0.001; 99% CI: –0.3651 to –0.2538) (**Fig. 4B**), indicating mRNAs of larger proteins tend to be depleted for AA-ending codons. The curve in **Fig. 4B** was steepest around 200 amino acids. Taking this as a cutoff, we found that of the 757 mRNAs encoding mature proteins ≤ 200 amino acids in length, 28 % (211) have an AA-ending codon usage rate ≥ 50% (**Fig. 4C**). For this group, there was a statistically significant, moderate negative correlation between AA % rates and corresponding mature protein sizes (*ρ* = -0.2590, two-tailed *p* < 0.001; 99% CI: –0.3466 to –0.1670) (**Fig. 4D**). In contrast, 3 % (33) of mRNAs encoding 1096 mature proteins in the 201 – 999 amino acid range have AA-ending codon rates ≥ 50% (**Fig. 4E**) and there was no statistically significant correlation between AA % rates and corresponding mature protein sizes for this group (*ρ* = 0.07054, two-tailed *p* = 0.02; 99% CI: –0.009560 to 0.1497) (**Fig. 4F**). Using the values in **Supplementary Table S4**, we calculated that for the 758 mRNAs encoding secreted mature proteins less than ≤ 200 amino acids in length, the average AA % rate was 34.4 %; for the 307 mRNAs in the secretome encoding mature proteins ≤ 100 amino acids in length, the average AA % rate was 39.45 %; and for the 64 mRNAs in the secretome encoding mature proteins ≤ 50 amino acids in length, the average AA % rate was 41.26 %. By comparison, for the 1854 genes in the secretome encoding mature proteins ≤ 999 amino acids in length, the average AA % rate was 26.16 %. We conclude that AA-ending codons, although rare in both the genome and the secretome compared to synonymous AG-ending codons, tend to accumulate in mRNAs encoding small, secreted proteins.

### *M. oryzae* mRNAs highly expressed *in planta* are depleted for AA-ending codons in favor of synonymous AG-ending codons

We hypothesized that mRNAs with high AA % rates in the secretome included those encoding uncharacterized cytoplasmic effectors, but which to validate experimentally? To understand how new cytoplasmic effectors may be more precisely identified, we sought to determine which uncharacterized, high AA % rate mRNAs – encoding small, secreted proteins – were expressed *in planta* during biotrophy and therefore likely to be physiologically relevant. We also sought more clarity on *M. oryzae* AA-ending codon usage during host infection. To achieve this, we took a proof-of-principle snapshot of the *in planta M. oryzae* transcriptome at 36 hpi. This is 12 hrs after cuticle penetration, when invasive hyphae (IH) are filling the first infected cell, all colonized rice cells are living, biotrophic interfaces are intact (Li et al., 2023b), and known biotrophic effector genes including *PWL2* and *BAS4* are being expressed and translated (**Fig. 1A**). We obtained RNA at 36 hpi, in triplicate, from three-week-old leaf sheaths of the susceptible rice line CO39 infected with Guy11. We performed RNAseq analysis and identified *M. oryzae* biotrophic growth-expressed genes using the 70-15 reference genome (**Supplementary Table S3**). We found that across the entire 36 hpi *M. oryzae in planta* biotrophic growth transcriptome, the AA-ending codon usage rate was 28.27 %, close to the genome average of 29 %, with only 5.7 % of detected genes having AA % rates ≥ 50 % (**Supplementary Table S5**). We normalized gene counts against *M. oryzae* actin expression, subtracted any values from mock infected control leaves, then ordered the resulting 10044 genes (that had both detectable expression and predicted protein coding regions) by relative normalized counts (**Fig. 5A**; values in **Supplementary Table S3**). A small number of *M. oryzae* genes were very highly expressed at 36 hpi, and the 28 most highly expressed genes with > 130 normalized counts (an arbitrary cut-off matching the steepest part of the curve in **Fig. 5A**) had an average AA % rate of 6.04 %, with five genes in this highly expressed set having 0 AA-ending codons (**Table 2**; **Supplementary Table S3**). Thus, very highly expressed *M. oryzae* biotrophic genes display a paucity of AA-ending codons. The genes in this highly expressed set were mostly associated with active growth processes and encoded ribosomal, translational, metabolic and cell-wall synthesis proteins, as exemplified by MGG_05247 (*GDH2*) (AA % rate = 12.65 %) encoding NAD-specific glutamate dehydrogenase (**Table 2**), a characterized gene essential for biotrophic growth (Li et al., 2023b). There were no secreted protein genes in this highly expressed group (**Table 2**). Furthermore, the 10 genes encoding ribosomal or translation-associated proteins with expression levels above the 130 count cut off have an average AA % rate of 4.6 % – for example, MGG_01742 encoding elongation factor 2 has 2 AA-and 142 AG-ending codons in 844 codons, yielding an AA % rate of 1.39 % (**Table 2**) – while in the top 100 most highly expressed genes, the 38 genes encoding putative ribosomal proteins have an average AA % rate of 3.51 % (**Supplementary Table S3**). These low AA % rates–indicating synonymous AG-ending codon-enrichment– are consistent with ribosomal proteins being efficiently translated from highly expressed mRNAs with optimal codons (Karlin and Mrazek, 2000; Mahlab and Linial, 2014) and are in sharp contrast to the high AA % rates for known cytoplasmic effector genes (*BAS1*, *AVR-Pik*, *AVR-Pita* and *PWL2*), encoding proteins experimentally determined to be secreted unconventionally into the BIC (Giraldo et al., 2013; Li et al., 2023a), which were expressed in our transcriptome at much lower levels (**Fig. 5A**). Two genes encoding experimentally determined apoplastic effectors, *BAS4* and *SLP1* (Li et al., 2023a), were expressed in our transcriptome at intermediate levels (**Supplementary Table S3**; **Fig. 5A**).

**Figure 5.**
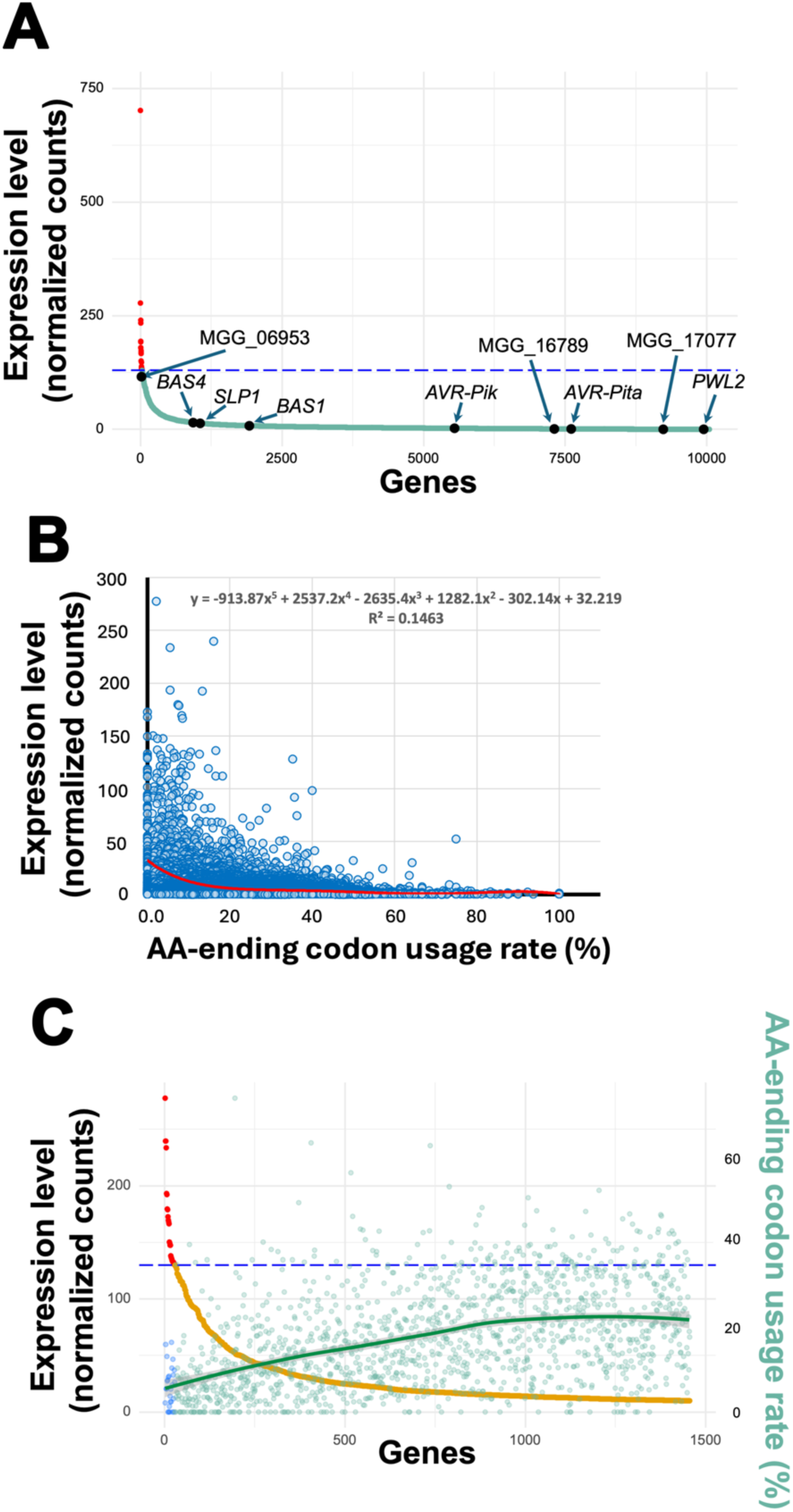
*M. oryzae* genes with low *in planta* expression accumulate AA-ending codons. **A**. Transcriptome data derived from RNAseq analysis of *M. oryzae* genes expressed during biotrophic growth in detached rice leaf sheaths at 36 hpi reveals few highly expressed genes and low-expressed known effector genes. 10044 genes were detected, and their expression levels were normalized. The horizontal dashed blue line marks an arbitrary threshold for high expression of 130 normalized counts. Red dots denote the 28 most highly expressed genes, while green dots represent genes with relatively lower expression. Nine selected genes are highlighted with black dots (see text for details). Values are the means of three biological replicates (**Supplementary Table S3**). **B**. At 36 hpi, AA % rates tend to increase in biotrophic transcriptome mRNAs as expression levels decrease, and there is thus a significant negative correlation between gene expression level and mRNA AA-ending codon usage rates. One outlier gene (MGG_03641) was excluded to improve curve fitting. The plot was generated using the scatter chart function in Microsoft Excel. The red curve represents a fifth-order polynomial regression fit (equation and R² shown). Spearman’s rank correlation analysis revealed a moderate but significant negative correlation between gene expression level and AA-ending codon rate (*ρ* = –0.2828, two-tailed *p* < 0.001; 99% CI: – 0.3070 to –0.2583). **C.** Normalized transcript counts were analyzed for 1454 genes, with a minimum expression threshold of 10 normalized counts. MGG_03641 was excluded as an outlier. Genes with normalized counts ≥ 130 are displayed as red dots, those with counts < 130 are shown as yellow dots. The blue dashed line represents the threshold value of 130. Blue dots on the far left represent AA % rates for the highly expressed genes with normalized counts ≥ 130, while green dots represent those with counts <130. The green trend line, showing increasing AA % rates with decreasing expression level, was generated using LOESS (locally estimated scatterplot smoothing) nonlinear regression to model the overall codon usage pattern.

**Table 2.** Highest expressed *M. oryzae* genes in the *in planta* transcriptome.

| Rank | Mean normalized counts | Protein name | Codons | #AA-ending | #AG-ending | AA % rate |
| --- | --- | --- | --- | --- | --- | --- |
| 1 | 701.9 | elongation factor 1-alpha | 473 | 3 | 84 | 3.45% |
| 2 | 277.8 | cell wall glucanosyltransferase | 367 | 1 | 45 | 2.17% |
| 3 | 239.78 | cross-pathway control protein 1 | 239 | 5 | 26 | 16.13% |
| 4 | 233.73 | ADP, ATP carrier protein | 315 | 2 | 34 | 5.56% |
| 5 | 193.5 | uncharacterized protein | 297 | 2 | 34 | 5.56% |
| 6 | 192.4 | amino-acid permease inda1 | 577 | 8 | 52 | 13.33% |
| 7 | 179.7 | plasma membrane ATPase | 926 | 11 | 138 | 7.38% |
| 8 | 178.9 | peptidyl-prolyl cis-trans isomerase | 215 | 2 | 24 | 7.69% |
| 9 | 172.7 | uncharacterized protein | 230 | 0 | 14 | 0.00% |
| 10 | 169.6 | ATP synthase subunit beta | 514 | 7 | 77 | 8.33% |
| 11 | 168.2 | 40S ribosomal protein S18 | 156 | 0 | 30 | 0.00% |
| 12 | 166.6 | formate dehydrogenase | 442 | 6 | 64 | 8.57% |
| 13 | 150.3 | elongation factor 2 | 844 | 2 | 142 | 1.39% |
| 14 | 149.5 | guanine nucleotide-binding protein | 316 | 0 | 39 | 0.00% |
| 15 | 147.7 | 60S acidic ribosomal protein P0 | 313 | 2 | 51 | 3.77% |
| 16 | 147.5 | 60S ribosomal protein L10a | 217 | 1 | 40 | 2.44% |
| 17 | 138.1 | ATP synthase subunit alpha | 551 | 5 | 86 | 5.49% |
| 18 | 137 | 60S ribosomal protein L3 | 392 | 2 | 79 | 2.47% |
| 19 | 136.1 | woronin body major protein | 417 | 12 | 60 | 16.67% |
| 20 | 135 | 60S ribosomal protein L2 | 254 | 4 | 34 | 10.53% |
| 21 | 133.8 | 40S ribosomal protein S3aE | 256 | 4 | 58 | 6.45% |
| 22 | 133.2 | uncharacterized protein | 150 | 0 | 11 | 0.00% |
| 23 | 133 | NAD-specific glutamate dehydrogenase | 1045 | 21 | 145 | 12.65% |
| 24 | 132.4 | 40S ribosomal protein S5 | 214 | 3 | 31 | 8.82% |
| 25 | 132.3 | heat shock protein SSB1 | 614 | 4 | 120 | 3.23% |
| 26 | 130.8 | uncharacterized protein | 229 | 0 | 24 | 0.00% |
| 27 | 130.7 | histone H3 | 136 | 3 | 27 | 10.00% |
| 28 | 130.5 | 60S ribosomal protein L33-B | 109 | 1 | 13 | 7.14% |

### High mRNA AA % rates correlate with low gene expression levels *in planta*

Highly expressed *M. oryzae* growth genes were low in AA % rates in our *in planta* transcriptome (**Table 2**), whereas known effectors were expressed at low levels (**Fig 5A**). We asked whether there was a correlation between AA % rates and gene expression level *in planta*. By applying Spearman correlation analysis to all detected genes in the transcriptome, we found a moderate but significant negative correlation between gene expression level and AA % rate (rate (*ρ* = –0.2828, two-tailed *p* < 0.001; 99% CI: –0.3070 to –0.2583) (**Fig. 5B**). We conclude that at 36 hpi, low-expressed mRNAs tend to carry more AA-ending codons than synonymous AG-ending codons, while highly expressed genes exhibit the reciprocal (**Fig. 5C**).

When considered together, our transcriptome results are in line with the notion that highly expressed genes have optimal codons, including AG-ending codons, that are recognized by abundant, non-thiolated tRNAs (Rezgui et al., 2013) with efficient codon-anticodon base pairing to enable rapid high-fidelity translation elongation, whereas rare codons that pause or slow elongation – allowing accurate co-translational folding of, for example, secreted and membrane proteins – accumulate in low expressed genes (Yu et al., 2015; Parvathy et al., 2021; Quax et al., 2015; Liu, 2020; Kudla and Plotkin, 2011; Mahlab and Linial, 2014; Chaney and Clark, 2015; Zhang and Qian, 2025). For the highly expressed genes in **Table 2**, where AG-ending codons occur 95.4 % of the time compared to AA-ending codons, these values strongly deviate from the codon usage average for AG-ending codons across the genome, which is 71 %.

### Two *bona fide* cytoplasmic effectors discovered by high mRNA AA % rates

We organized the transcriptome in **Supplementary Table S3** by AA % rates (**Supplementary Table S5**). Genes with the highest AA % rates had very low expression levels (**Table 3**). 7 transcripts had 100 %, AA % rates, but two of these encoded putative membrane proteins (MGG_02805 and MGG_15126) and the others lacked SP-encoding sequences. 6 of the 7 were not enriched for AA-ending codons. The genes with the next 5 highest AA % rates, from 93.75 % to 87.88 %, included *AVR-PIK* (MGG_15972) (92 % AA) and *AVR-PITA* (MGG_15370) (87.88 % AA), MGG_16514 encoding a likely membrane bound protein, and two small (mature protein <100 amino acids), uncharacterized secreted proteins: MGG_17077 (93.75 % AA, 15 AA- and 1 AG-ending codons in 113 codons) and MGG_16789 (90 % AA, 9 AA- and 1 AG-ending codons in 90 codons) (**Table 3**; **Fig 5A**; **Supplementary Table S5**). In the secretome, MGG_17077 and MGG_16789 lie either side of *AVR-Pik* with regards to AA % rate (**Fig. 6A**), but they are uncharacterized (although they are not members of the sequence-unrelated, structurally homologous MAX effector family) (Seong and Krasileva, 2019). BLAST analysis showed they share no identity with each other or with *AVR-Pik* despite having similar AA % rates. The transcriptome data suggested they both encode proteins physiologically relevant to the infection process, therefore we proceeded to validate their localization by live-cell imaging. We replaced the *PWL2* coding sequence in pBV591 with the gene encoded either at MGG_17077 (**Fig. 6B**) or MGG_16789 (**Fig. 6C**) and expressed these genes fused to *mCherry:NLS*, under the *PWL2* promoter, in the WT Guy11 and Δ*uba4* mutant strains; these strains also expressed *BAS4* fused to *eGFP* under its own promoter. The proteins encoded at both MGG_17077 and MGG_16789 localized to the BIC in WT, indicating they are *bona fide* cytoplasmic effectors, and their production was dependent on Uba4. BFA treatment internalized Bas4-eGFP in both WT strains but BIC accumulation of the proteins encoded at MGG_17077 and MGG_16789 was unaffected (**Figs. 6B,C**), indicating the MGG_17077 and MGG_16789 effectors are secreted into the BIC in an unconventional manner. Treating Δ*uba4* infected rice cells with paromomycin remediated MGG_17077 and MGG_16789 protein production in this mutant strain (**Figs. 6B,C**), indicating AA-ending codons in the associated mRNAs require Uba4 for efficient decoding. Together, we conclude that the proteins encoded at MGG_17077 and MGG_16789 are small cytoplasmic effectors secreted into the BIC via the Type IV Golgi bypass UPS pathway. Moreover, their low-expressed, AA-ending codon-enriched mRNAs are subject to Uba4 translation control.

**Figure 6.**
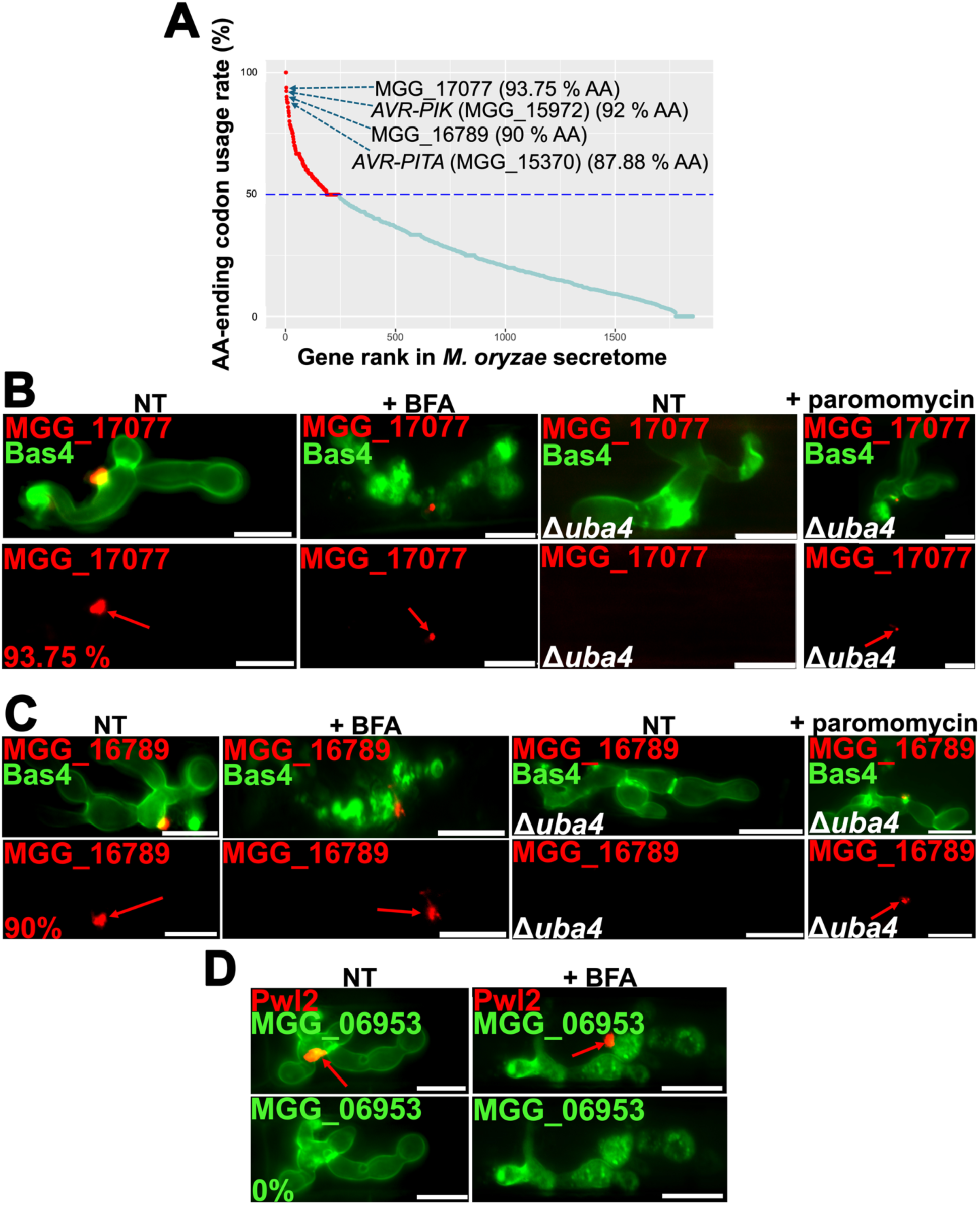
AA % rates coupled with *in planta* mRNA expression levels uncovers new *M. oryzae* effectors. **A**. MGG_17077 and MGG_16789 mRNAs have some of the highest AA % rates in the secretome. The 1854 genes in the *M. oryzae* secretome were ranked in descending order based on the AA % rate of each gene. The blue dashed line indicates 50 % AA rate. Red dots are genes with ≥ 50 % AA rates, green dots are genes with < 50 % AA rates. **B-D**. Live-cell images of the indicated strains and treatments were taken at 36 hpi for the non-treated (NT) controls. For BFA treatments, samples were treated at 32 hpi and imaged at 37 hpi. For paromomycin treatments, samples were treated at 36 hpi and imaged at 44 hpi. Red arrows indicate effector accumulation in the BIC. Bars are 10 µm. % value refers to the AA % rate for the indicated effector mRNA (Li et al., 2023a). Images are representative of 50 infected rice cell observations per leaf sheath, repeated in triplicate.

**Table 3.** mRNAs with the highest AA % rates in the *M. oryzae in planta* transcriptome.

| Rank | Mean normalized counts | Gene | Codons | #AA-ending | #AG-ending | AA % rate |
| --- | --- | --- | --- | --- | --- | --- |
| 1 | 1.10167112 | MGG_02805 | 64 | 1 | 0 | 100.00% |
| 2 | 0.2528569 | MGG_15126 | 102 | 2 | 0 | 100.00% |
| 3 | 0.22292377 | MGG_16144 | 60 | 4 | 0 | 100.00% |
| 4 | 0.15432084 | MGG_15066 | 64 | 4 | 0 | 100.00% |
| 5 | 0.11176687 | MGG_15529 | 84 | 1 | 0 | 100.00% |
| 6 | 0.11135873 | MGG_16609 | 79 | 1 | 0 | 100.00% |
| 7 | 0.11054243 | MGG_14909 | 64 | 11 | 0 | 100.00% |
| 8 | 0.18241505 | MGG_17077 | 113 | 15 | 1 | 93.75% |
| 9 | 2.13946925 | *AVR-Pik* | 113 | 12 | 1 | 92.31% |
| 10 | 0.81603697 | MGG_16789 | 90 | 9 | 1 | 90.00% |
| 11 | 0.12499768 | MGG_16514 | 127 | 9 | 1 | 90.00% |
| 12 | 0.7297091 | *AVR-Pita* | 224 | 29 | 4 | 87.88% |

### Discovery of a new apoplastic effector by coupling low mRNA AA % rates with high *in planta* expression levels

In contrast to the low expression, high AA % rates of MGG_17077 and MGG_16789, the highest expressed gene from the secretome detected in our transcriptome, and the 29^th^ most highly expressed biotrophic gene at this 36 hpi snapshot, was MGG_06953 (**Fig. 5A**; **Supplementary Table S3**) encoding a putative uncharacterized protein with a 0 % AA rate (0 AA- and 13 AG-ending codons in 138 codons; mature protein size = 123 aa). We predicted this gene would encode an apoplastic effector. We exchanged the *BAS4* promoter and coding region in pBV591 with the gene encoded at the MGG_06953 allele to express *MGG_06953* fused to *eGFP* under the *RP27* promoter (along with *PWL2-mCherry:NLS* under its native promoter) in WT. As expected, the protein encoded at MGG_06953 accumulated in the apoplast (**Fig 6D**). Treatment with BFA retained the MGG_06953 protein in IH, indicating it is secreted into the apoplast via the conventional ER-Golgi pathway (**Fig 6D**). We conclude that the gene at MGG_06953 encodes a new apoplastic effector, highly expressed during invasive biotrophic growth.

### Limited distribution of the new effectors across host-infecting lineages

BLAST analysis of the protein coding sequences showed orthologues of the apoplastic effector-encoding gene MGG_06953 from 70-15 are present in a Brazilian isolate of the related species *Pyricularia pennisetigena* (PpBr36_07430), in *P. grisea* (PgNI_06037), and it overlaps with a gene in the *M. oryzae* Zambian wheat blast isolate ZM2-1 (on CP099696.1). A copy of the 70-15 cytoplasmic effector-encoding gene MGG_17077 is present in the *M. oryzae* Zambian wheat blast isolate ZM2-1 (on CP099699.1). The 70-15 cytoplasmic effector-encoding gene MGG_16789 has an orthologue in the *M. oryzae* Zambian wheat blast isolate ZM2-1 (on CP099698.1) and shares identify with the *P. grisea* gene PgNI_09257. We conclude that these effectors are not widespread in *M. oryzae* lineages and thus may contribute to host species specificity, but some occur in related *Pyricularia* species.

## Discussion

Plant pathogens mediate infection by deploying host-specific effector proteins with roles in inhibiting or evading plant innate immunity. Characterizing effector action and their host targets aids the dissection of plant innate immunity (Toruño et al., 2016), while cytoplasmic effectors can be used to find or design cognate R proteins (Maidment et al., 2023), together enhancing disease resistance breeding (Wilson and McDowell, 2022). The bottleneck in fungi, however, is the paucity of known cytoplasmic effectors due to an incomplete understanding of effector deployment and evolution. Effector classes are poorly conserved at the nucleotide level and difficult to deduce from first principles in fungal genomes. This is because although all effector genes carry signal peptide sequences that can be identified computationally, and while apoplastic and cytoplasmic effectors are secreted by different routes, there are no known motifs or sorting signals specifying the unconventional secretion of cytoplasmic effectors into the BIC and their targeting to the host cell. Thus, effectors are difficult to classify computationally, and cytoplasmic effector discovery is hampered. Here, by assessing AA-ending codon usage in mRNAs encoding SP-carrying proteins, we correctly identified novel *bona fide* cytoplasmic effectors in an unbiased manner by computational means.

As proof-of-principle, we showed that mRNA AA-ending codon usage rates, coupled with a snapshot of mRNA abundance during biotrophy, accurately distinguished cytoplasmic from apoplastic effectors, a major advance in our tool set for effector discovery. Two cytoplasmic effectors were successfully predicted from high AA % rates and low *in planta* transcript abundance (which were also features of other known cytoplasmic effector mRNAs detected in our transcriptome), and a new apoplastic effector was predicted from its high *in planta* expression level and 0 % AA rate. Both new cytoplasmic effectors could have been identified in the secretome from their high AA % rates alone, but their detection in the *in planta* transcriptome gave confidence that they were physiologically relevant. Earlier or later transcriptomes may prioritize other candidates. Also, in other fungi where the secretome is not defined, infection-related transcriptomics may find new cytoplasmic effectors based on low expression levels and high AA % rates.

Although effectors are often sequence-unrelated, recently, genome-wide computational structural biology approaches using template-free modeling have clustered effectors into structurally analogous groups based on their predicted protein folds (Seong and Krasileva, 2021; Yan et al. 2023). Previously, we showed that known *M. oryzae* MAX family cytoplasmic effectors carrying the sequence-unrelated six-strand β-sandwich fold were enriched for rare AA-ending codons, suggesting they are under dual selection for both the sequence-unrelated protein fold and for codon usage (Li et al., 2023a). For the cytoplasmic effectors identified in this study, structural analysis indicates that the effector encoded at MGG_17077 carries a predicted glycosyl hydrolase fold, and the effector encoded at MGG_16789 has a cystatin-like domain; both folds are also carried by other predicted *M. oryzae* effectors (Seong and Krasileva, 2021). Therefore, these new cytoplasmic effectors are from two different sequence-unrelated, structurally analogous groups (Seong and Krasileva, 2021), suggesting dual selection for both sequence-unrelated protein folds and AA-ending codon-enrichment is widespread in cytoplasmic effector mRNAs. The nature of this dual selection is unclear, but the non-neutral selection for AA-ending codons may lie in co-translational folding requirements for cytoplasmic effector peptides destined for the Golgi bypass UPS pathway, which could differ from those of ER-Golgi secreted apoplastic effectors (Li et al., 2023a). AA-ending codons are slowly decoded by thiolated tRNAs to limit translation elongation rates and allow accurate co-translational folding (Chaney and Clark, 2015; Collart and Weiss, 2019; Parvathy et al., 2022). Slow elongation of translation using non-optimal AA-ending codons matched to low abundant tRNAs can prevent peptide misfolding and reduce the frequency of ribosome stalling resulting from ribosome collisions (Novoa and Pouplana, 2012). This notion of AA-ending codons contributing to accurate folding of cytoplasmic effectors in *M. oryzae* is supported by the converse finding that the most highly expressed growth-related genes in our 36 hpi snapshot, encoding cellular proteins, tended to carry optimal AG-ending codons matching abundant tRNAs with few or no rare AA-ending codons matched to less abundant tRNAs, thereby ensuring fast translation. Apart from the highly expressed MGG_06953 gene, apoplastic effector mRNAs tended to be intermediate in their expression levels and AA-ending codon usage. Taken together, we conclude that rare AA-ending codons are overemphasized in low-expressed cytoplasmic effector genes from across taxa, suggesting that their accumulation in this context is strongly non-neutral (Zhang and Qian, 2025) and is in addition to – and possibly independent of – selection for sequence-unrelated, structurally analogous effector folds likely required for effector function.

Where tested, the decoding of rare AA-ending codons in *M. oryzae* cytoplasmic effector mRNAs requires Uba4-dependent tRNA thiolation, which may provide another layer of translation elongation precision. In addition to folding accuracy, the precise modulation of translation elongation may allow nascent peptides to interact with specific binding partners (Gasparski et al., 2023), for example UPS pathway-specific SNAREs (Dulal and Wilson, 2024), to target them to the correct secretion pathway.

tRNA reusage could be an alternative or additional explanation for the high AA-ending codon rates in cytoplasmic effector mRNAs. Codons recognizing the same tRNA isoacceptor, even rare codons, are often overrepresented in a gene (Cannarozzi et al., 2010); having this enrichment may optimize genes for tRNA reuse, raising the local concentration of tRNAs and leading to efficient translation (Cannarozzi et al., 2010; Novoa and de Pouplana, 2012). If tRNA recycling is more efficient for smaller mRNAs, this could account for the observed trend that *M. oryzae* AA % rates increased with decreasing mature protein size. Two possible reasons why tRNA recycling may be important for cytoplasmic effector mRNA translation is that firstly, biotrophic growth occurs under nutrient-limiting conditions (Li et al., 2023b), while at the same time, cytoplasmic effectors must be synthesized promptly and deployed into the host to suppress immunity. Accumulating codons infrequently used by the rest of the genome in cytoplasmic effector mRNAs could generate a pool of recycled cognate tRNAs not in demand by more highly expressed genes, thereby reducing competition for tRNAs and optimizing their translation. This would be consistent with the observation that genes requiring optimal expression under starvation response conditions are enriched in rare codons (Elf et al., 2003; Cannarozzi et al., 2010). Secondly, it is conceivable that proteins secreted by the UPS pathway are synthesized in different microcompartments under different cellular conditions compared to those destined for the ER-Golgi pathway, hence the dependance of the former on both tRNA thiolation and, potentially, a local pool of otherwise low abundance tRNAs. Future work may distinguish these scenarios and clarify the link between tRNA modification and tRNA reusage, if any.

The AA % rate rule operates across *M. oryzae* host infecting-lineages and closely related species. Currently, only *M. oryzae* (Giraldo et al., 2013) and the oomycete *P. infestans* (Wang et al., 2017) have been shown experimentally to secrete cytoplasmic and apoplastic effectors by different pathways, with the experimentally confirmed SP-carrying *P. infestans* cytoplasmic effector gene *Pi04313* having a higher AA % rate (44 %) than the confirmed *P. infestans* apoplastic effector gene *EPIC1* (14 %) (Dulal and Wilson, 2024). Genes in other fungal or oomycete secretomes with elevated AA % rates may therefore indicate the similar use of an unconventional secretion pathway, permitting comparisons that may help us to understand why such pathways are utilized for host cell-deployed effectors. Future work may also determine how the effectors encoded at MGG_06953, MGG_17077 and MGG_16789 act during infection, as well as revisit Avr-Pi54 to understand how an apoplast-localized protein interacts with its cognate host R protein. The correct prediction of Avr-Pi54 as an apoplast-localized effector based solely on mRNA AA % rates lends considerable support to our notion that AA % rates are unbiased predictors of effector deployment routes. Future challenges will involve determining whether there is a clear transition in AA % rates between cytoplasmic and apoplastic effector mRNAs and if so, finding where that boundary lies. If there is instead an overlap between cytoplasmic and apoplastic effector mRNA AA % rates at moderate values, as suggested by the continuous curve when secretome genes are plotted against AA % rates, then experimentally determined expression levels and mature protein size will be important for separating them.

Together, this work sheds new light on cytoplasmic effector properties and evolution and takes us a step closer to understanding how and why cytoplasmic effectors are discriminated from apoplastic effectors during secretion. AA-ending codon enrichment is likely a fundamental principle of cytoplasmic effector biology, and our findings will aid the search for new effectors across taxa, which is expected to bolster food security. This work also provides more information on the connection between AA-ending codon usage and unconventional protein secretion, a poorly understood role for codon usage bias in any system (Li et al., 2023a).

## Data Availability

The data discussed in this publication have been deposited in NCBI’s Gene Expression Omnibus (Edgar et al., 2002) and are accessible through GEO Series accession number GSE297821 (https://www.ncbi.nlm.nih.gov/geo/query/acc.cgi?acc=GSE297821). The strains and oligonucleotides used in this study are listed in **Supplementary Tables S1** and **S2**. The gene sequences used in the course of this study can be found at NCBI under the following accession numbers: MGG_13863 for *PWL2*, MGG_10914 for *BAS4*, MT669814 for *PWL1*, LC202655 for *PWT4*, AB498874 for *Avr-Pii*, MN072523 for *AvrPib*, MGG_12655 for *AvrPi9*, HF545677 for *AVR-Pi54* and MGG_17077, MGG_16789 and MGG_06953 for the new effector genes characterized in this study. The algorithms used are described in the methods section and are publicly accessible on GitHub.

## Author contributions

R.A.W. conceived and supervised the project and acquired funding. G.L. performed the bioinformatic analyses. S.T., N.D. and B.W. performed the molecular and cell biology experiments. G.L., S.T., N.D. and R.A.W. analyzed the data. R.A.W. wrote the manuscript. All authors reviewed and edited the manuscript.

## Materials and methods

### Fungal strains and culture conditions

The fungal strains used and generated during the course of this study are listed in **Supplementary Table S1**. The *M. oryzae* wild-type (WT) rice isolate Guy11 (Wilson and Talbot, 2009) and the Δ*uba4* mutant strain derived from Guy11 (Li et al., 2023a) were used as the parental strains. All strains are stored as filter stocks at -20 °C. For downstream applications, three colonies of each strain were grown on a single plate of complete media (Li et al., 2015) and spores were harvested at 3 – 5 days for all *UBA4^+^*strains and at 7 – 10 days for all Δ*uba4* strains.

### Vector and strain construction

New vectors used in this study were constructed at GenScript USA, Inc., using the vector pBV591 (Khang et al., 2010) as the base template (Li et al., 2023a). Briefly, the *PWL2* sequence and native promoter were replaced in pBV591 by *PWL1* or *PWT4* and their respective promoters using protein coding sequences (CDSs) obtained from NCBI. The *PWL2* sequence alone was replaced in pBV591 by the *AVR-Pi54* CDS obtained from NCBI. The *PWL2* sequence alone was replaced in pBV591 by the CDSs encoded at the MGG_17077 and MGG_16879 alleles, which were obtained from the *M. oryzae* genome sequence database at Ensembl Fungi (https://fungi.ensembl.org/Magnaporthe_oryzae/Info/Index). The *BAS4* CDS and promoter was replaced in pBV591 by the CDS at MGG_06953, which was obtained from Ensembl Fungi, and by the *RP27* promoter sequence (Richter et al., 2024), respectively. The indicated genes were synthesized at GenScript USA, Inc., cloned onto their respective vectors using the Clone EZ cloning method, and confirmed by sequencing. The vectors were transformed into Guy11 and Δ*uba4* protoplasts generated as described previously (Li et al., 2015). Transformants were initially selected on the basis of hygromycin B (Neta Scientific, Inc., USA) resistance, then confirmed by PCR; the primers used for screening transformants are in **Supplementary Table S2**. The PCR-confirmed transformants were tested for fluorescence on detached rice sheaths until a minimum of three independent transformants per vector per strain had been identified and shown to be identical with regards to effector production and deployment. One transformant per vector per strain was used for downstream analyses.

### Live-cell imaging

For live-cell imaging, following harvesting, spores were washed in ddH_2_O and resuspended in 2 ml 0.2 % gelatin (Acros, USA), counted on a hemocytometer, adjusted to a concentration of 2 × 10^5^ spores ml^−1^, and applied to optically clear detached rice leaf sheaths from 3.5 -4-week-old seedlings of the susceptible rice cultivar CO-39. Each strain and treatment was tested on leaf sheaths from at least three different plants. Detached rice leaf sheaths infected with the indicated strains were incubated in all cases at 26 °C in the dark under high humidity (75%). Images were obtained at 32-36 hr post inoculation (hpi), unless otherwise stated, using a Nikon Eclipse Ni-E upright microscope and NIS Elements software.

Excitation/emission was 488 nm/505–531 nm for eGFP and 543 nm/590–632 nm for mCherry. For paromomycin treatments, fresh 1 mg ml^−1^ paromomycin (Sigma-Aldrich, USA) was prepared in 0.2 % gelatin, added to cells at 36 hpi, and the effects observed at 44 hpi. For the Brefeldin A (BFA) (Fisher Scientific, USA) assays, 4-week-old rice sheaths were inoculated with 2 × 10^5^ conidia ml^−1^ of each strain. At 32 hpi, the infected leaf sheath samples were trimmed into ultrathin sections and placed into 60 mm petri-dishes containing either freshly prepared BFA (50 µg ml^−1^ in 0.1 % DMSO in H_2_O) or a mock control (0.1% DMSO in H_2_O) and left for 5 hours with agitation at 23 °C. Then, the sections were mounted on slides using forceps and imaged with the Nikon Eclipse Ni-E Upright Microscope. Representative images are based on the observations of 50 infected rice cells per leaf sheath per treatment, repeated in triplicate, as standard (Li et al., 2023a; Li et al., 2023b).

### tRNA gene counts

The number of tRNA genes in the 70-15 genome that match AA- or AG-ending codons was retrieved from the Genomic tRNA Database (https://gtrnadb.org) from predictions made by tRNAscan-SE (Chan et al., 2021).

### RNA extraction and RNAseq analysis

Detached leaf sheaths from 4-week-old rice plants were inoculated with spores of the *M. oryzae* wild-type strain Guy11 and suspended in 0.2 % gelatin at a concentration of 1 X 10⁵ spores ml^−1^. Leaf sheaths treated with mock solution (0.2 % gelatin) were the control. At 36 hpi, all leaf sheaths were harvested and flash-frozen in liquid nitrogen as a single batch over a 15-minute period. Three biological replicates per treatment were collected for subsequent RNAseq analysis. To extract RNA, leaf sheath tissues were ground in liquid nitrogen, and total RNA was extracted using the TRIzol reagent (Invitrogen, USA), according to the manufacturer’s protocol. After removing contaminating genomic DNA using DNase I, total RNA was purified with the PureLink RNA Mini kit (Thermo Fisher Scientific, USA), following the manufacture’s instructions. Total RNA was run on a denaturing agarose gel to estimate quality, and RNA quantity was calculated using a Nanodrop Spectrophotometer. RNA integrity and purity were assessed using the Agilent Technologies 2100 Bioanalyzer, and high quality (RIN ≥ 7.0) was confirmed for all samples. For each sample, 3 μg of total RNA at a concentration of more than 100 ng/μl was used in for library construction.

The Poly(A) RNA sequencing libraries were prepared following Illumina’s TruSeq-stranded-mRNA sample preparation protocol. mRNAs with poly(A) tails were isolated from total RNA using oligo-(dT) magnetic beads and two rounds of purification. The purified poly(A) mRNA fractions were cleaved into smaller fragments using a divalent cation buffer at elevated temperatures. The cleaved RNA fragments were reverse-transcribed to create the final cDNA library in accordance with a strand-specific library preparation by dUTP method. Quality control analysis and quantification of the sequencing library were performed using Agilent Technologies 2100 Bioanalyzer High Sensitivity DNA Chip. The average insert size for the paired-end libraries was 300 ± 50 bp. cDNA library sequencing was performed with 150-nucleotide pair-end reads on Illumina’s NovaSeq 6000 sequencing system at LC Sciences (USA) following the vendor’s recommended protocol. Three biological replications were used for RNAseq experiments.

For the quantification of gene expression in Guy11 during biotrophic growth at 36 hpi, raw sequencing data were first filtered by removing contaminated adaptor sequences, low quality bases and undetermined bases using CUTADAPT (Martin, 2011) and perl scripts in LC Science. Quality control of raw sequencing data was performed using FastQC (Andrews, 2010), and summary reports were generated with MultiQC (Ewels et al., 2016). 9 bp of low-quality bases from the 5’-ends of the reads were removed using SEQTK (version 1.2, (https://github.com/lh3/seqtk), and high quality valid reads were mapped to the *Magnaporthe oryzae* reference genome (https://ftp.ncbi.nlm.nih.gov/genomes/all/GCF/000/002/495/GCF_000002495.2_MG8/GCF_000002495.2_ MG8_genomic.fna.gz) using SUBREAD for Unix (version 2.0; Liao et al., 2013). The alignment was carried out for read groups of all samples, including all replications of the Guy11-infected leaf sheaths and the mock control. Gene annotation as GTF (version 2.2, https://ftp.ncbi.nlm.nih.gov/genomes/all/GCF/000/002/495/GCF_000002495.2_MG8/GCF_000002495.2_ MG8_genomic.gtf.gz) was obtained from NCBI, corresponding to the MG8 genome assembly GCF_000002495.2. Read counts per-gene were computed using FEATURECOUNTS (Liao et al., 2014) from SUBREAD (version 2.0; Liao et al., 2013). Counts per million (CPM) values were used as normalized expression levels for each gene. To avoid normalization bias introduced by mock-inoculated samples and to ensure proper incorporation of sequencing depth, library sizes were set based on total tissue read counts as determined by FEATURECOUNTS. Genes with CPM > 0 in at least two samples were retained for further analysis. Raw counts of the selected genes were then used to identify differentially expressed genes (DEGs) using Quasi-likelihood methods (Lund et al., 2012) implemented in edgeR (version 4.4.2, Robinson et al., 2010). Differential expression was assessed by comparing gene expression in samples inoculated with the wild-type strain Guy11 versus mock controls, with DEGs defined as those exhibiting a false discovery rate (FDR) < 0.05 and an absolute log_2_ fold change (log_2_FC) ≥ 1.

Using this approach, a total of 10044 *M. oryzae* genes were identified as expressed in Guy11-infected rice leaf sheath tissues. All count data and edgeR analysis results are provided in **Supplementary Table S3**. For each gene in this dataset, functional and pathway annotations were obtained through interactive data mining using the BioMart (http://www.biomart.org/, version 0.7), Gene Ontology (http://geneontology.org/, Release 2022-01-13), and KEGG (https://www.genome.jp/kegg/, Release 101.0, January 1, 2022) databases. These annotations were subsequently integrated in R (version 4.1.2).

Additionally, due to gaps in the aforementioned databases, annotation information for several genes was manually retrieved from NCBI and incorporated accordingly.

### AA-ending codon rate analysis

For codon usage analysis, the *Magnaporthe oryzae* MG8 coding sequence (CDS) database (*Magnaporthe_oryzae.MG8.cds.all.fa.gz*, https://ftp.ensemblgenomes.ebi.ac.uk/pub/fungi/release-58/fasta/magnaporthe_oryzae/cds/) was used as the reference CDS dataset for *M. oryzae* genes. Codon usage patterns were analyzed using the open-source R package coRdon (version 1.24.0; https://www.bioconductor.org/packages/release/bioc/html/coRdon.html; Elek et al., 2024). For each gene, the AA-ending codon usage rate was calculated as the number of AA-ending codons in the CDS divided by the total number of AA-ending and AG-ending codons, multiplied by 100. AA-ending codons include AAA, CAA, and GAA, while AG-ending codons include AAG, CAG, and GAG.

For codon usage analysis of the *Magnaporthe oryzae* secretome, a reference dataset comprising 1854 predicted secreted proteins, < 1000 amino acids in length, was used. These proteins were identified previously by Seong and Krasileva (Seong and Krasileva, 2021).

### Data correlation analysis and plot generation

During the correlation analysis, data normality was first assessed using Gaussian distribution analysis in PASW Statistics 18.0 (PASW Statistics Inc.). For datasets that did not meet the assumption of normality, correlations were evaluated using the Spearman rank correlation method in GraphPad Prism 8. To visualize the correlations, non-linear regression analysis was performed in Microsoft Excel to generate the regression equation and curve. The correlation coefficient (*ρ*, rho) and two-tailed *p*-value were calculated and used to determine the statistical significance of the correlations, with a threshold of *p* < 0.05 indicating significance. Negative correlations were interpreted based on the value of *ρ* (rho), ranging from -0.1 to -1.0. Specifically: no correlation: *ρ* between 0.0 and -0.1; weak negative correlation: *ρ* between -0.1 and -0.25; moderate negative correlation: *ρ* between -0.25 and -0.5; strong negative correlation: *ρ* between -0.5 and -1.0. LOESS analysis was performed following the method of Cleveland and Devlin (Cleveland and Devlin 1988).

All other plots generated from bioinformatics analysis were generated using ggplot2 (version 3.5.2; Wickham, 2009) in R (version 4.4.0). Bioinformatics analyses performed following the adaptor removal utilized the high-performance computing resources provided by the Holland Computing Center, University of Nebraska-Lincoln, USA. All in-house scripts used for differential gene expression analyses, AA-ending codon analyses, and figure generation are available on GitHub: https://github.com/gangli/novel_Cytoplasmic_Effectors/tree/main, and archived on Zenodo at DOI: https://doi.org/10.5281/zenodo.15469270

### Statistical analysis

In addition to the statistical approaches mentioned above, average AA % rates were calculated from individual AA % rates in the *M. oryzae* secretome using Microsoft Excel.

## References

Andrews S. (2010) FastQC: a quality control tool for high throughput sequence data. http://www.bioinformatics.babraham.ac.uk/projects/fastqc.

Cannarozzi G, et al., A role for codon order in translation dynamics. Cell. 2010; 141: 355–367. doi: 10.1016/j.cell.2010.02.036.

Chan PP, Lin BY, Mak AJ and Lowe TM. tRNAscan-SE 2.0: improved detection and functional classification of transfer RNA genes. Nucleic Acids Res. 2021; 49: 9077–9096. doi: 10.1093/nar/gkab688.

Chaney JL and Clark PL. Roles for synonymous codon usage in protein biogenesis. Annu Rev Biophys. 2015; 44: 143–66. doi: 10.1146/annurev-biophys-060414-034333.

Cleveland WS and Devlin SJ. Locally-weighted regression: an approach to regression analysis by local fitting. J Am Statl Assoc. 1988; 83: 596–610.

Collart MA and Weiss B. Ribosome pausing, a dangerous necessity for co-translational events. Nucleic Acids Res. 2020; 48:1043–1055. doi: 10.1093/nar/gkz763.

Dean RA, et al., The genome sequence of the rice blast fungus *Magnaporthe grisea*. Nature. 2005; 434: 980–986. doi: 10.1038/nature03449.

Dulal N and Wilson RA. Paths of least resistance: Unconventional effector secretion by fungal and oomycete plant pathogens. Mol Plant Microbe Interact. 2024; 37: 653–661. doi: 10.1094/MPMI-12-23-0212-CR.

Edgar R, Domrachev M and Lash AE. Gene Expression Omnibus: NCBI gene expression and hybridization array data repository. Nucleic Acids Res. 2002; 30: 207–210. doi: 10.1093/nar/30.1.207.

Elek A, Kuzman M and Vlahovicek K. (2024) coRdon: Codon sage analysis and prediction of gene expressivity. R package version 1.24.0, https://github.com/BioinfoHR/coRdon.

Elf J, Nilsson D, Tenson T and Ehrenberg M. Selective charging of tRNA isoacceptors explains patterns of codon usage. Science. 2003; 300: 1718–1722. doi: 10.1126/science.1083811.

Ewels P, Magnusson M, Lundin S and Käller M. MultiQC: summarize analysis results for multiple tools and samples in a single report. Bioinformatics. 2016; 32: 3047–3048. doi: 10.1093/bioinformatics/btw354.

Gasparski AN, Moissoglu K, Pallikkuth S, Meydan S, Guydosh NR and Mili S. mRNA location and translation rate determine protein targeting to dual destinations. Mol Cell. 2023; 83: 2726–2738.e9. doi: 10.1016/j.molcel.2023.06.036.

Giraldo MC, et al., Two distinct secretion systems facilitate tissue invasion by the rice blast fungus *Magnaporthe oryzae*. Nat Commun. 2013; 4: 1996. doi: 10.1038/ncomms2996.

Goffena J, et al., Elongator and codon bias regulate protein levels in mammalian peripheral neurons. Nat Commun. 2018; 9: 889. doi: 10.1038/s41467-018-03221-z.

Gómez Luciano LB, et al., Blast fungal genomes show frequent chromosomal changes, gene gains and losses, and effector gene Ttrnover. Mol Biol Evol. 2019; 36: 1148–1161. doi: 10.1093/molbev/msz045.

Inoue Y, et al., Evolution of the wheat blast fungus through functional losses in a host specificity determinant. Science. 2017; 357: 80–83. doi: 10.1126/science.aam9654.

Jones JD and Dangl JL. The plant immune system. Nature. 2006; 444: 323–329. doi: 10.1038/nature05286.

Karlin S and Mrázek J. Predicted highly expressed genes of diverse prokaryotic genomes. J Bacteriol. 2000; 182: 5238–5250. doi: 10.1128/JB.182.18.5238-5250.2000.

Khang CH, et al., Translocation of *Magnaporthe oryzae* effectors into rice cells and their subsequent cell-to-cell movement. Plant Cell. 2010; 22: 1388–1403. doi: 10.1105/tpc.109.069666.

Kourelis J, Marchal C, Posbeyikian A, Harant A and Kamoun S. NLR immune receptor-nanobody fusions confer plant disease resistance. Science. 2023; 379: 934–939. doi: 10.1126/science.abn4116.

Li G, Marroquin-Guzman M and Wilson RA. Chromatin Immunoprecipitation (ChIP) Assay for Detecting Direct and Indirect Protein -DNA Interactions in Magnaporthe oryzae. Bioprotocol. 2015; 5: e1643. doi: 10.21769/BioProtoc.1643.

Li G, Dulal N, Gong Z and Wilson RA. Unconventional secretion of *Magnaporthe oryzae* effectors in rice cells is regulated by tRNA modification and codon usage control. Nat Microbiol. 2023a; 8:1706–1716. doi: 10.1038/s41564-023-01443-6

Li G, Gong Z, Dulal N, Marroquin-Guzman M, Rocha RO, Richter M and Wilson RA. A protein kinase coordinates cycles of autophagy and glutaminolysis in invasive hyphae of the fungus *Magnaporthe oryzae* within rice cells. Nat Commun. 2023b; 14: 4146. doi: 10.1038/s41467-023-39880-w.

Liao,Y, Smyth G. and Shi W. The Subread aligner: fast, accurate and scalable read mapping by seed-and-vote. Nucleic Acids Res. 2013; 41: e108. doi: 10.1093/nar/gkt214.

Liao Y, Smyth GK and Shi W. featureCounts: an efficient general purpose program for assigning sequence reads to genomic features. Bioinformatics. 2014; 30: 923–930. doi: 10.1093/bioinformatics/btt656.

Liu Y. A code within the genetic code: codon usage regulates co-translational protein folding. Cell Commun Signal. 2020; 18: 145. doi: 10.1186/s12964-020-00642-6.

Lund SP, Nettleton D, McCarthy DJ and Smyth GK. Detecting differential expression in RNA-sequence data using quasi-likelihood with shrunken dispersion estimates. Stat Appl Genet Mol Biol. 2012; 11: Article 8. doi: 10.1515/1544-6115.1826.

Mahlab S and Linial M. Speed controls in translating secretory proteins in eukaryotes–an evolutionary perspective. PLoS Comput Biol. 2014; 10: e1003294. doi: 10.1371/journal.pcbi.1003294.

Maidment JHR, et al., Effector target-guided engineering of an integrated domain expands the disease resistance profile of a rice NLR immune receptor. Elife. 2023; 12: e81123. doi: 10.7554/eLife.81123.

Martin M. Cutadapt removes adapter sequences from high-throughput sequencing reads. EMBnet.journal. 2011; 17: 10–12. doi: 10.14806/ej.17.1.200.

Masaki HI, et al., Host specificity controlled by *PWL1* and *PWL2* effector genes in the finger millet blast pathogen *Magnaporthe oryzae* in eastern Africa. Mol Plant Microbe Interact. 2023; 36: 584–591. doi: 10.1094/MPMI-01-23-0012-R.

Nedialkova DD and Leidel SA. Optimization of codon translation rates via tRNA modifications maintains proteome integrity. Cell. 2015; 161: 1606–1618. doi: 10.1016/j.cell.2015.05.022.

Novoa EM and Ribas de Pouplana L. Speeding with control: codon usage, tRNAs, and ribosomes. Trends Genet. 2012; 28: 574–581. doi: 10.1016/j.tig.2012.07.006.

Novoa EM, Pavon-Eternod M, Pan T and Ribas de Pouplana L. A role for tRNA modifications in genome structure and codon usage. Cell. 2012; 149: 202–213. doi: 10.1016/j.cell.2012.01.050.

Oliveira-Garcia E, et al., Clathrin-mediated endocytosis facilitates the internalization of *Magnaporthe oryzae* effectors into rice cells. Plant Cell. 2023; 35: 2527–2551. doi: 10.1093/plcell/koad094.

Parvathy ST, Udayasuriyan V and Bhadana V. Codon usage bias. Mol Biol Rep. 2022; 49: 539–565. doi: 10.1007/s11033-021-06749-4

Plotkin JB and Kudla G. Synonymous but not the same: the causes and consequences of codon bias. Nat Rev Genet. 2011; 12: 32–42. doi: 10.1038/nrg2899.

Quax TE, Claassens NJ, Söll D and van der Oost J. Codon bias as a means to fine-tune gene expression. Mol Cell. 2015; 59: 149–161. doi: 10.1016/j.molcel.2015.05.035.

Ray S, et al., Analysis of *Magnaporthe oryzae* genome reveals a fungal effector, which is able to induce resistance response in transgenic rice line containing resistance gene, *Pi54*. Front Plant Sci. 2016; 7: 1140. doi: 10.3389/fpls.2016.01140.

Rezgui VA, et al., tRNA tK^UUU^, tQ^UUG^, and tEUUC wobble position modifications fine-tune protein translation by promoting ribosome A-site binding. Proc Natl Acad Sci U S A. 2013; 110: 12289–12294. doi: 10.1073/pnas.1300781110.

Richter M, et al., Membrane fluidity control by the *Magnaporthe oryzae* acyl-CoA binding protein sets the thermal range for host rice cell colonization. PLOS Pathog. 2024; 20: e1012738. doi: 10.1371/journal.ppat.1012738.

Robinson MD, McCarthy DJ. and Smyth GK. edgeR: A Bioconductor package for differential expression analysis of digital gene expression data. Bioinformatics. 2010; 26: 139–140. doi: 10.1093/bioinformatics/btp616.

Seong K and Krasileva KV. Computational structural genomics unravels common folds and novel families in the secretome of fungal phytopathogen *Magnaporthe oryzae*. Mol Plant Microbe Interact. 2021; 34: 1267–1280. doi: 10.1094/MPMI-03-21-0071-R

Sperschneider J and Dodds PN. EffectorP 3.0: Prediction of apoplastic and cytoplasmic effectors in fungi and oomycetes. Mol Plant Microbe Interact. 2022; 35: 146–156. doi: 10.1094/MPMI-08-21-0201-R.

Toruño TY, Stergiopoulos I and Coaker G. Plant-pathogen effectors: Cellular probes interfering with plant defenses in spatial and temporal manners. Annu Rev Phytopathol. 2016; 54: 419–441. doi: 10.1146/annurev-phyto-080615-100204.

Tuller T, et al., An evolutionarily conserved mechanism for controlling the efficiency of protein translation. Cell. 2010; 141: 344–354. doi: 10.1016/j.cell.2010.03.031.

Valent B. Dynamic gene-for-gene interactions undermine durable resistance. Mol Plant Microbe Interact. 2025; 38: 104–117. doi: 10.1094/MPMI-02-25-0022-HH.

Wang S, Boevink PC, Welsh L, Zhang R, Whisson SC and Birch PRJ. Delivery of cytoplasmic and apoplastic effectors from *Phytophthora infestans* haustoria by distinct secretion pathways. New Phytol. 2017; 216: 205–215. doi: 10.1111/nph.14696.

Wickham, H. ggplot2: Elegant graphics for data analysis. Springer, New York, 2009

Wilson RA and Talbot NJ. Under pressure: investigating the biology of plant infection by *Magnaporthe oryzae*. Nat Rev Microbiol. 2009; 7:185–95. doi: 10.1038/nrmicro2032.

Wilson RA and McDowell JM. Recent advances in understanding of fungal and oomycete effectors. Curr Opin Plant Biol. 2022; 68: 102228. doi: 10.1016/j.pbi.2022.102228.

Wu J, et al., Comparative genomics identifies the *Magnaporthe oryzae* avirulence effector *AvrPi9* that triggers *Pi9*-mediated blast resistance in rice. New Phytol. 2015; 206: 1463–1475. doi: 10.1111/nph.13310.

Yan X, et al. The transcriptional landscape of plant infection by the rice blast fungus *Magnaporthe oryzae* reveals distinct families of temporally co-regulated and structurally conserved effectors. Plant Cell. 2023; 35: 1360–1385. doi: 10.1093/plcell/koad036.

Yu CH, et al., Codon usage influences the local rate of translation elongation to regulate co-translational protein folding. Mol Cell. 2015; 59: 744–754. doi: 10.1016/j.molcel.2015.07.018.

Zhang J and Qian W. Functional synonymous mutations and their evolutionary consequences. Nat Rev Genet. 2025; 26: 789–804. doi: 10.1038/s41576-025-00850-1.

Zhang X, et al., tRNA thiolation optimizes appressorium-mediated infection by enhancing codon-specific translation in *Magnaporthe oryzae*. Nucleic Acids Res. 2025; 53: gkae1302. doi: 10.1093/nar/gkae1302.

