## Supplementary Table S1. for "AA-ending codon-based computational analyses uncover novel cytoplasmic effectors in the *Magnaporthe oryzae* secretome"

### Supplementary Table S1. Strains used in this study.

| **Strains** | **Genotype** | **Reference** |
| --- | --- | --- |
| Guy11 | *M. oryzae* wild type isolate (WT) used throughout this study | Wilson and Talbot, 2009 |
| Δ*uba4* | WT parental strain carrying a deletion of the *UBA4* gene (MGG_05569) encoding Urm1-activating enzyme | Li et al. 2023a |
| *PWL2-mCherry:NLS*  *BAS4-GFP* | WT Guy11 strain carrying pBV591 (Khang et al. 2010) expressing *PWL2* fused to *mCherry:NLS* under its native promoter and *BAS4* fused to *eGFP* under its native promoter | Li et al. 2023a |
| Δ*uba4*  *PWL2-mCherry:NLS*  *BAS4-GFP* | Δ*uba4* strain carrying pBV591 (Khang et al. 2010) expressing *PWL2* fused to *mCherry:NLS* under its native promoter and *BAS4* fused to *eGFP* under its native promoter | Li et al. 2023a |
| *PWL1-mCherry:NLS*  *BAS4-GFP* | WT Guy11 strain carrying a modified pBV591 vector expressing the *PWL1* protein coding sequence fused to *mCherry:NLS* under its native promoter and *BAS4* fused to *eGFP* under its native promoter | *This study* |
| Δ*uba4*  *PWL1-mCherry:NLS*  *BAS4-GFP* | Δ*uba4* strain carrying a modified pBV591 vector expressing the *PWL1* protein coding sequence fused to *mCherry:NLS* under its native promoter and *BAS4* fused to *eGFP* under its native promoter | *This study* |
| *PWT4-mCherry:NLS*  *BAS4-GFP* | WT Guy11 strain carrying a modified pBV591 vector expressing the *PWT4* protein coding sequence fused to *mCherry:NLS* under its native promoter and *BAS4* fused to *eGFP* under its native promoter | *This study* |
| Δ*uba4*  *PWT4-mCherry:NLS*  *BAS4-GFP* | Δ*uba4* strain carrying a modified pBV591 vector expressing the *PWT4* protein coding sequence fused to *mCherry:NLS* under its native promoter and *BAS4* fused to *eGFP* under its native promoter | *This study* |
| *AVR-Pi54-mCherry:NLS*  *BAS4-GFP* | WT Guy11 strain carrying a modified pBV591 vector expressing the *AVR-Pi54* protein coding sequence fused to *mCherry:NLS* under the *PWL2* promoter and *BAS4* fused to *eGFP* under its native promoter | *This study* |
| Δ*uba4*  *AVR-Pi54-mCherry:NLS*  *BAS4-GFP* | Δ*uba4* strain carrying a modified pBV591 vector expressing the *AVR-Pi54* protein coding sequence fused to *mCherry:NLS* under the *PWL2* promoter and *BAS4* fused to *eGFP* under its native promoter | *This study* |
| *MGG_17077-mCherry:NLS*  *BAS4-GFP* | WT Guy11 strain carrying a modified pBV591 vector expressing the protein coding sequence at MGG_17077 fused to *mCherry:NLS* under the *PWL2* promoter and *BAS4* fused to *eGFP* under its native promoter | *This study* |
| Δ*uba4*  *MGG_17077-mCherry:NLS*  *BAS4-GFP* | Δ*uba4* strain carrying a modified pBV591 vector expressing the protein coding sequence at MGG_17077 fused to *mCherry:NLS* under the *PWL2* promoter and *BAS4* fused to *eGFP* under its native promoter | *This study* |
| *MGG_16789-mCherry:NLS*  *BAS4-GFP* | WT Guy11 strain carrying a modified pBV591 vector expressing the protein coding sequence at MGG_16789 fused to *mCherry:NLS* under the *PWL2* promoter and *BAS4* fused to *eGFP* under its native promoter | *This study* |
| Δ*uba4*  *MGG_16789-mCherry:NLS*  *BAS4-GFP* | Δ*uba4* strain carrying a modified pBV591 vector expressing the protein coding sequence at MGG_16789 fused to *mCherry:NLS* under the *PWL2* promoter and *BAS4* fused to *eGFP* under its native promoter | *This study* |
| *MGG_06953-mCherry:NLS*  *BAS4-GFP* | WT Guy11 strain carrying a modified pBV591 vector expressing the protein coding sequence at MGG_06953 fused to *mCherry:NLS* under the *RP27* promoter and *BAS4* fused to *eGFP* under its native promoter | *This study* |
| Δ*uba4*  *MGG_06953-mCherry:NLS*  *BAS4-GFP* | Δ*uba4* strain carrying a modified pBV591 vector expressing the protein coding sequence at MGG_06953 fused to *mCherry:NLS* under the *RP27* promoter and *BAS4* fused to *eGFP* under its native promoter | *This study* |

Khang CH, et al. Translocation of *Magnaporthe oryzae* effectors into rice cells and their subsequent cell-to-cell movement. *Plant Cell*. 2010; 22: 1388-1403.

Li G, Dulal N, Gong Z and Wilson RA. Unconventional secretion of *Magnaporthe oryzae* effectors in rice cells is regulated by tRNA modification and codon usage control. *Nat. Microbiol*. 2023a; 8:1706-1716.

Wilson RA and Talbot NJ. Under pressure: investigating the biology of plant infection by *Magnaporthe oryzae*. *Nat. Rev. Microbiol*. 2009; 7:185-95.
