## Supplementary Table S2. for "AA-ending codon-based computational analyses uncover novel cytoplasmic effectors in the *Magnaporthe oryzae* secretome"

**Supplementary Table S2. Oligonucleotides used in this study.**

| **Primer Name** | **Sequence** |
| --- | --- |
| Avr-Pi54_Conf Forward | GAGCTCAAGGCCGTCTTC |
| GFP_Conf -Reverse | TGTTGTGGCGGATCTTGAA |
| PWl2 Conf Fw | CTTATTATGGTCCCGGGTGAT |
| qGFP Re | TGTTGTGGCGGATCTTGAA |
| qBas4 Re | GCATCCGAATGGCAGAGT |
| RP27 Fw | CACGTTGTCGCCTAACAGAT |
| qGFPRe | TGTTGTGGCGGATCTTGAA |
